# Stereotypical interciliary contacts in a *C. elegans* sense organ

**DOI:** 10.64898/2026.04.10.717756

**Authors:** Nikhila Krishnan, Samantha Leslie, Hannah Lawson, Yu-Ming Lu, Leland Wexler, Kirsten Judge, Maxwell G. Heiman, Piali Sengupta

## Abstract

Physical interactions among cells and their processes are critical for intercellular communication and the generation of ordered tissue patterns. Primary cilia projecting from the cell surface have recently been shown to form contacts with the processes of diverse cell types, as well as with other cilia, in the brain and other organs. Whether these ciliary contacts are established in an instructive manner or are formed passively due to physical proximity is unclear. Ultrastructural analyses previously showed that the cilia of a subset of sensory neurons in the head amphid organs of *C. elegans* exhibit interciliary contacts within a glia-defined channel. Here we show that these ciliary contact patterns are stereotyped and can be established in the absence of neighboring cilia, indicating that these associations may not simply reflect relative positioning within the amphid channel. We show that mutations in genes implicated in ciliary protein trafficking, ciliary membrane phospholipid composition, and cilia-cell interactions disrupt cilia structure and/or interciliary contacts, and that in a subset of mutants, cilia with altered morphologies can nevertheless establish correct contacts. Together, our findings suggest that cilia-cilia interactions within a sense organ are established via instructive mechanisms, and raise the possibility that cellular functions may be modulated by cilia-mediated intercellular communication.

**Summary:** This work investigates how primary cilia, structures that detect and transmit signals, form contacts with one another in a head sensory organ of the nematode *Caenorhabditis elegans*. The authors found that these contacts follow consistent patterns and can form even when neighboring cilia are absent, suggesting they are actively established rather than occurring due to physical proximity. The authors identified mutations in genes that regulate cilia protein content, membrane composition, and cellular adhesion that disrupt cilia structure or contacts. These findings suggest that cilia-cilia interactions are regulated, raising the possibility that they play important roles in cell communication.

## INTRODUCTION

Precise orchestration of interactions among cells and cellular processes during development allows for the emergence of ordered patterns. Stereotypy in these patterns across scales is essential for coordinating the properties of cells, tissues, and organs, thereby enabling coherent functional output. A key feature underlying these organizational patterns is selective cell-cell interactions dictated by developmentally programmed expression of molecules that mediate cell adhesion or repulsion via homophilic or heterophilic interactions (Gumbiner 1996; Steinberg and Takeichi 1994; Townes and Holftreter 1955). The critical importance of regulated cellular interactions is particularly evident in the nervous system whose function relies on the establishment of ordered neuronal connections with target cells (Agi et al. 2020). The selection of a specific wiring partner among a set of possible targets is thought to be driven by the combinatorial expression of specific cell adhesion molecules in each cell type leading to the establishment of defined connections (Sanes and Zipursky 2020; Sperry 1963; Togashi et al. 2009). However, it has also been suggested that neurons can form connections indiscriminately with multiple adjacent partners, suggesting that physical proximity can instruct connectivity (Cook et al. 2023; Packer et al. 2013; Peters and Feldman 1976; Rees et al. 2017).

Primary cilia are small microtubule-based organelles that project from the surface of nearly all cell types including neurons in the central nervous system of metazoans. These structures house a range of signal transduction molecules and play a critical role in transmitting environmental information to the cell (Jurisch-Yaksi et al. 2024; Singla and Reiter 2006). The location and orientation of a cilium vary by cell and tissue type. Sensory neurons contain specialized cilia at their dendritic ends that are exposed to the environment, whereas in central neurons, cilia typically extend from the soma and exhibit defined orientations in specific brain regions (Falk et al. 2015; Jurisch-Yaksi et al. 2024; Monfared et al. 2025; Ott et al. 2024; Wu et al. 2024). Recent ultrastructural analyses have shown that neuronal and glial cilia in the brain form extensive and unique sets of contacts with dendrites, axons and glial processes, and are also found in close apposition to synaptic clefts (Ott et al. 2024; Sheu et al. 2022; Wu et al. 2024). Similarly, cilia of pancreatic beta cells contact islet cells, axons of innervating neurons, and cilia of other beta cells (Muller et al. 2024). Interciliary contacts have also been observed in rod photoreceptors, cholangiocytes, and cultured MDCK cells (Ott et al. 2012). It is currently unknown how stereotyped these ciliary contacts are *in vivo*, and whether these contacts are formed via instructive mechanisms or are established passively with neighboring cells and/or cellular processes.

Only sensory neurons in *C. elegans* are ciliated, and as in other organisms, cilia are present at their dendritic ends and are exposed directly or indirectly to the external or internal environment (Doroquez et al. 2014; Perkins et al. 1986; Ward et al. 1975). Twelve pairs of ciliated sensory neurons are present in the bilateral head amphid sense organs; each of these neuron pairs responds to a unique set of external stimuli to drive behavior (Ferkey et al. 2021; Goodman and Sengupta 2019). Cilia of eight of these amphid sensory neurons are located in a channel (“channel cilia”) comprised of the processes of the amphid sheath (AMsh) and socket (AMso) glial cells (Fig. 1a), whereas the cilia of the remaining four neurons are embedded within these glial processes (Doroquez et al. 2014; Perkins et al. 1986; Ward et al. 1975). Using transmission electron microscopy (TEM), we previously showed that channel cilia appear to be organized in a stereotypical manner in each of the bilateral amphid channels, and that distal ciliary segments contact each other (Doroquez et al. 2014; Ward et al. 1975). The extent of this stereotypy, and whether this organization arises as a consequence of actively regulated mechanisms is unclear.

**Fig. 1.**
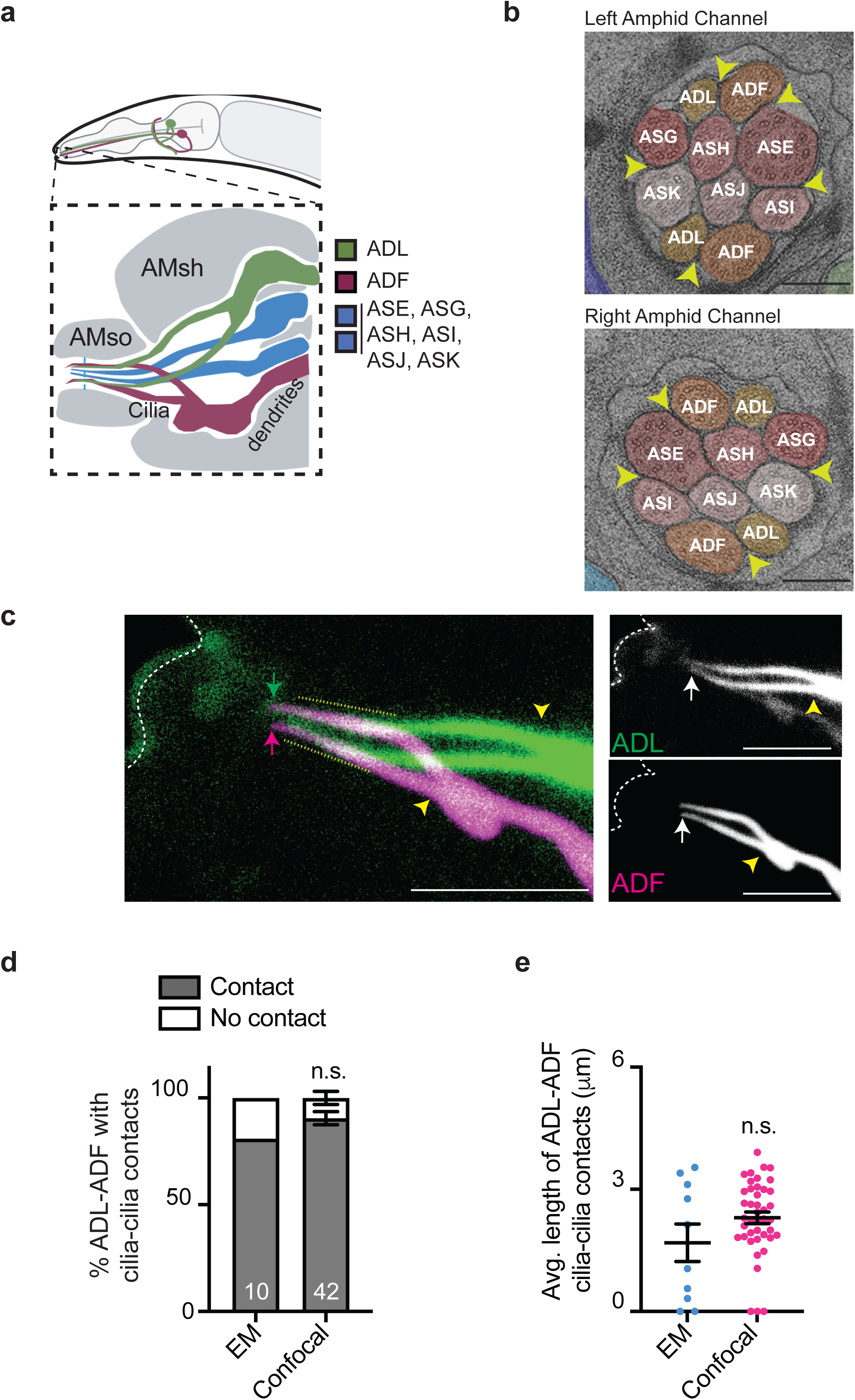
Distal segments of channel neuron cilia establish a stereotyped organization and pattern of interciliary contacts. **a)** Cartoon showing cilia of a subset of sensory neurons in the amphid channel. Only two of the six monociliated neurons are shown (blue). AMsh: amphid sheath process; AMso: amphid socket process. Anterior at left. **b)** Cross-section TEM image of the left and right amphid channels at the approximate level indicated by a dotted blue line in **a**. Cilia of each channel neuron are labeled. Arrowheads indicate electron densities at interciliary contacts. Scale bars: 250 nm. Adapted from (Doroquez et al. 2014). **c)** Representative maximum intensity projection images of ADL and ADF cilia visualized via expression of *srh-220*p*::gfp* (ADL) and *srh-142*p*::dsRed* (ADF). Dashed white line: worm nose; arrows: cilia tip; yellow arrowheads: cilia base. Yellow dotted lines: extent of interciliary contacts. Anterior at left. Scale bars: 5 μm. **d)** Percentage of ADL and ADF neuron pairs examined via TEM or confocal microscopy exhibiting contacts between cilia (see Methods). Numbers indicate the number of neuron pairs examined. Confocal data are from three biologically independent experiments. n.s.: not significant (Fisher’s exact test). **e)** Extent of interciliary contacts between ADL and ADF cilia (see Methods) as assessed via TEM or confocal microscopy. Each circle is the measurement from a single pair of neurons from one amphid channel; n = 10 and 42 neuron pairs for the TEM and confocal data, respectively. Confocal data are from three biologically independent experiments. Horizontal and vertical bars: Mean and SEM. n.s.: not significant (Mann-Whitney test). *Alt text*: Images and quantification of interciliary contacts between the cilia of the ADL and ADF neurons in the amphid sense organ of *C. elegans*.

Here we show that channel cilia exhibit an invariant pattern of organization and interciliary contacts in the adult *C. elegans* hermaphrodite. We find that individual and pairs of cilia can elongate in the amphid channel and establish their pattern of interciliary contacts even in the absence of other cilia in the channel. We show that mutations in a subset of ciliary genes disrupt cilia morphology and/or interciliary contacts, but that cilia with severely altered structures can nevertheless establish the correct interciliary contacts. Our results imply that cilia organization in the channel may be established by a regulated process possibly via cilia-localized adhesion molecules.

## MATERIALS and METHODS

### C. elegans strains

All *C. elegans* strains were maintained on plates containing Nematode Growth Medium (NGM) agar seeded with *E. coli* OP50 at 15°C or 20°C. The presence of mutations in strains was verified by PCR-based amplification and/or sequencing. Experimental plasmids were injected at 10-25 ng/μl along with an *unc-122*p::fluorescent reporter as the coinjection marker at 40-50 ng/μl. One day old well-fed adult hermaphrodites were used for imaging studies. Strains used in this study are listed in Supplementary Table 1.

### Molecular biology

Plasmids were generated using traditional cloning methods. A construct containing the mNeptune coding sequence was a kind gift from Eviatar Yemini. The *osm-3* mutant phenotype was rescued via expression of an *osm-3(oy156ts)* cDNA under cell-specific promoters; this mutation does not disrupt OSM-3 function at permissive temperatures (15°C-25°C) (Philbrook et al. 2024). All plasmids were verified by PCR and/or sequencing. Plasmids used in this study are listed in Supplementary Table 2.

### Electron microscopy measurements

IMOD Open Source software from the University of Colorado was used to quantify interciliary contacts shown in Fig. 1 (Kremer et al. 1996). Quantification was performed using transmission electron microscopy data reported previously (Doroquez et al. 2014). To measure the extent of ADL-ADF ciliary overlap, ADL and ADF cilia were identified by tracing individual channel cilia distally through serial sections to their transition zones in order to identify these biciliated cells. The number of sections in which interciliary contacts were detectable were identified and counted. The final number of sections were multiplied by the thickness of each serial ultrathin section (70 nm) to obtain the total interciliary contact length.

### Light microscopy

Growth-synchronized one-day old well-fed adult hermaphrodites were anaesthetized on 1% agarose pads containing 10mM Tetramisole Hydrochloride (Sigma Aldrich L9756) prior to imaging. Imaging of ADL and ADF cilia in Fig. 1 was performed using a Zeiss LSM 880 laser scanning confocal inverted microscope. Zeiss Zen software (black edition) was used to acquire the images with *z*-step sizes of 0.2 μm using a 60x oil objective with an optical zoom of 10x. A subset of imaging was performed on an inverted Zeiss Axiovert with a Yokogawa CSU-X1 spinning disk confocal unit and a Photometrics Quantum 512SC camera. Slidebook 6.0 software was used to acquire images with *z*-step sizes of 0.2 μm using a 100x oil immersion objective. Additional imaging data was acquired on a DM6000 Leica inverted microscope with a CSU-W1 spinning disk confocal unit and a ZL41 scMOS camera. Fusion 2.4.0.14 software was used to acquire images with *z*-step sizes of 0.2 μm using a 100x oil objective. Control and experimental strains were imaged in parallel using the same conditions in individual experiments unless indicated otherwise. Images were analyzed using FIJI/ImageJ (NIH) and Imaris (Oxford Instruments). Measurements were performed using the Fiji/ImageJ segmented line tool for all line-based measurements unless specified otherwise. All image analyses were performed on maximum intensity projections from confocal micrographs.

### Image analyses

#### Cilia length

Cilia length was measured from the distal tip of the cilium (arrows in images) to the cilia base (yellow arrows in images).

#### Location of cilia base

The distance between the bases of ADL and ADF cilia was measured between their periciliary membrane compartments (PCMCs) (see Fig. 4a). The location of each cilia base relative to the worm nose was measured by quantifying the distance between the worm nose tip visualized by autofluorescence to the cilia base (see Fig. 4a).

#### Interciliary contact

Cilia length and cilia overlap were quantified for each pair of neurons within an animal. The maximum number of interciliary contacts between the two cilia each of ADF and ADL in each animal is 4; that between ASE and ASH or ASH and ASI is 2. Since not all cilia were visible in each *z*-stack, the number of interciliary contacts possible was adjusted to the total number of cilia that were observed. The percentage of cilia exhibiting interciliary contacts was quantified as the number of cilia expressing GFP whose fluorescence overlapped with that of cilia expressing a red fluorescent marker divided by the total number of possible interciliary contacts. The extent of interciliary overlap between ADL and ADF was quantified as the average length of fluorescence overlap between each of the cilia in a pair of ADL and ADF neurons (see Fig. 1c). Since a subset of cilia showed length differences across mutant and rescue lines, the extent of interciliary overlap was normalized to cilia length: average length of interciliary contacts/(neuron 1 cilia length + neuron 2 cilia length). The length of two cilia was averaged per neuron for ADL or ADF.

ASH and ASI cilia lie in close apposition in the distal amphid channel making them difficult to resolve by light microscopy in laterally oriented animals, as the cilia of these neurons stack along the optical axis. Therefore, we selected animals positioned with either their dorsal or ventral side facing the coverslip. This orientation allowed both amphid channels to be imaged with the cilia positioned approximately in parallel and located in closely adjacent *z*-planes. For consistency, contacts between ASH and ASE cilia were also examined in animals with the same orientation.

#### Disorganized vs organized cilia

‘Collapsed’ cilia were defined as those in which the two cilia each of ADL or ADF contact each other either throughout their length or at their distal ends (representative examples are shown in Supplementary Fig. 2). All other morphology defects in these cilia were quantified under the “other” category.

### Statistical analyses

All graphs were generated and statistical analyses performed using GraphPad Prism 10. All shown data are from a minimum of two biologically independent experiments; the exact number of independent experiments is indicated in each Figure legend. Statistical tests used and the number of experimental samples are also described in each Figure legend. Since a subset of data did not meet normality assumptions, statistical significance was determined using the non-parametric Mann-Whitney test, or Kruskal-Wallis test with Dunn’s correction for multiple comparisons. For binned data, statistical significance was assessed using Fisher’s exact test with Holm’s correction for multiple comparisons.

## RESULTS

### A subset of sensory neuron cilia in the amphid sense organs exhibits stereotypical contacts

Since we previously examined only six amphid channels via TEM (Doroquez et al. 2014), we used light microscopy to further assess the stereotypy of the observed ciliary contacts across larger sample sizes (Turan et al. 2025). Six channel neurons contain a single cilium each, whereas the ADL and ADF sensory neurons are biciliated (Doroquez et al. 2014; Perkins et al. 1986; Ward et al. 1975) (Fig. 1a). Although the ADF and ADL dendrites enter the amphid pore at distinct locations, the cilia of these neurons are in close apposition in the distal channel, with a cilium of one neuron contacting a cilium of the second neuron on either side of the channel (Fig. 1b) (Doroquez et al. 2014). The membranes of adjacent cilia are within 12 nm as assessed via TEM (Fig. 1b). We noted electron densities between ADF and ADL cilia (16/20 contacts between ADL and ADF cilia exhibited densities; n=5 animals) as well as at a subset of contacts between other cilia pairs (Fig. 1b), raising the possibility that these cilia may be connected via gap or adhesion junctions (Wu et al. 2024). We focused our attention primarily on the biciliated ADF and ADL neurons.

As shown in Fig. 1c, while the bases of ADF and ADL cilia are located at a distance of ∼4 μm from each other in the proximal channel, the distal segments of ADF and ADL cilia marked via the expression of fluorescent reporters exhibited extensive overlap. As also observed via TEM, a cilium of one neuron contacted a cilium of the second neuron (Doroquez et al. 2014) (Fig. 1c, Supplementary Video 1). ∼90% of amphid channels examined via light microcopy showed a similar pattern of ADF and ADL cilia contacts (Fig. 1d). Moreover, the extent of overlap between individual cilia pairs (see Methods) was consistent between analyses performed via TEM and light microscopy (Fig. 1e). We infer that ADF and ADL cilia exhibit a stereotyped pattern of contacts in the distal amphid channel.

### Cilia of biciliated ADL and ADF neurons can partly establish contacts in the absence of distal ciliary segments of other amphid channel neurons

ADF and ADL cilia not only contact each other but also contact cilia of other channel neurons at their distal segments (Fig. 1b) (Doroquez et al. 2014; Ward et al. 1975). The contacts between ADL and ADF cilia may arise passively due to the spatial organization of cilia within the amphid channel or may be regulated by specific interciliary adhesion mechanisms. To distinguish between these hypotheses, we examined whether contacts between ADF and ADL cilia can be established in the absence of other channel cilia.

Middle segments of channel cilia are built by the partly redundant functions of the heterotrimeric kinesin-II and homodimeric OSM-3 IFT motors, whereas the distal segments are built by OSM-3 alone (Perkins et al. 1986; Signor et al. 1999; Snow et al. 2004) (Fig. 2a). Thus, channel cilia are fully truncated in animals doubly mutant for the *kap-1* subunit of kinesin-II and *osm-3*, whereas only the distal segments are truncated in *osm-3* mutants alone. As expected, cilia of both ADF and ADL were severely shortened in *kap-1; osm-3* double mutants with concomitant loss of interciliary contacts (Supplementary Fig. 1a-c). Expression of functional *osm-3* sequences specifically in ADL and ADF in the kinesin double mutant background lengthened the cilia of either neuron to only a minor extent (Supplementary Fig. 1a-c), indicating that OSM-3 alone is not sufficient to grow these cilia, or that these cilia are unable to elongate in the absence of other cilia in the channel. However, since both ADF and ADL cilia extend significantly upon shifting adult animals doubly mutant for a *kap-1* null allele and a temperature-sensitive (*ts*) allele of *osm-3* (Philbrook et al. 2024) to the permissive temperature (Judge et al, 2026) (Supplementary Fig. 1d-e), we infer that OSM-3-driven extension of these cilia in the *kap-1; osm-3* null mutant background requires the presence of one or more channel cilia.

**Fig. 2.**
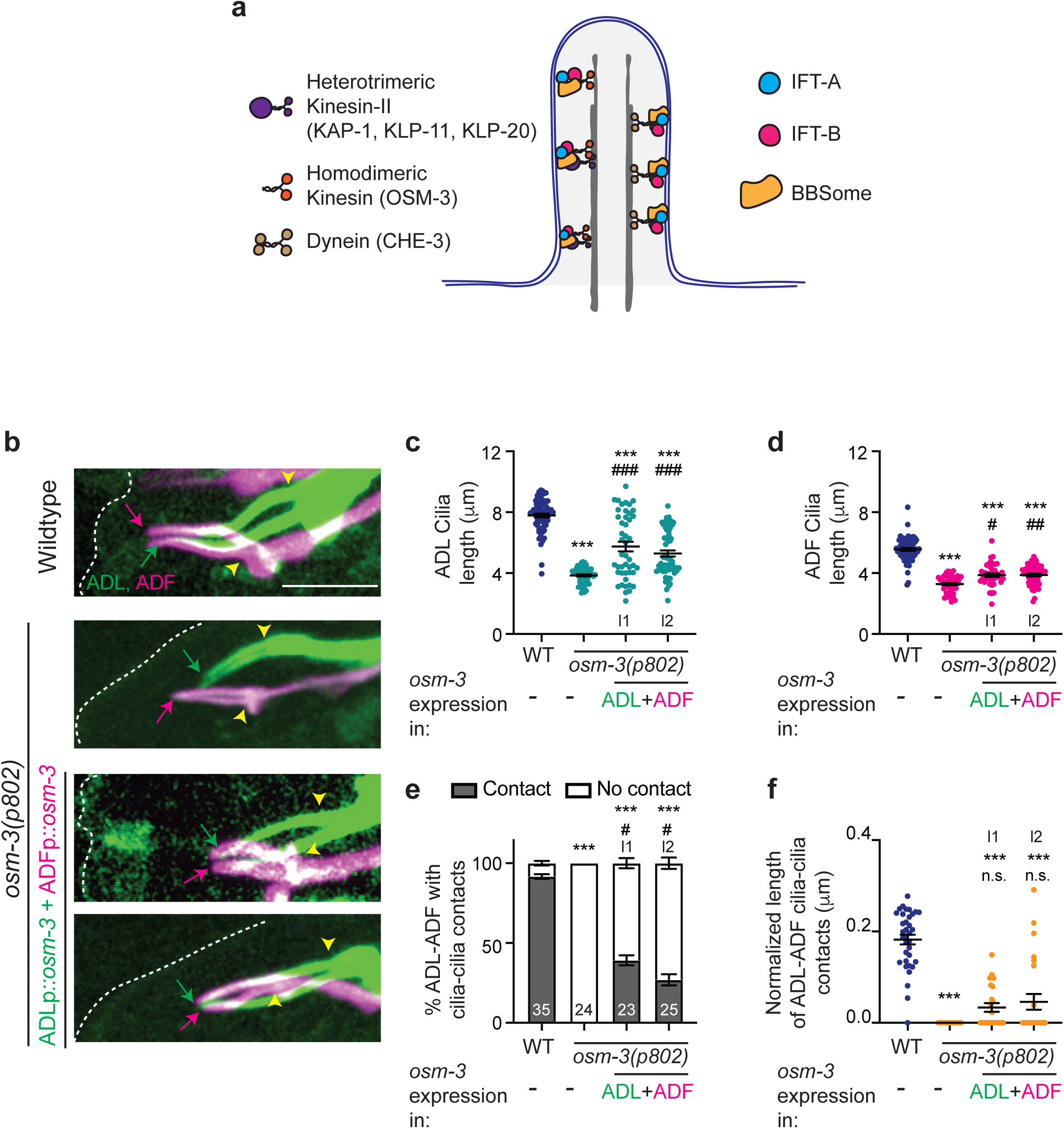
Interciliary contacts between ADF and ADL can be formed in the absence of distal segments of other amphid channel cilia. **a)** Cartoon of a model channel cilium in *C. elegans.* The middle ciliary segment is built by the kinesin-II and OSM-3 motors, whereas the distal segments are built by OSM-3 alone. **b)** Representative maximum intensity projection images of ADL and ADF cilia in the indicated genetic backgrounds. Dashed white line: worm nose; arrows: cilia tip; yellow arrowheads: cilia base. Anterior at left. Scale bar (for all images): 5 μm. **c,d)** Length of ADL (c) and ADF cilia in animals with the indicated genetic backgrounds. Each circle is the length of a single cilium of each neuron. n>46 neurons for ADL and ADF, n>44 neurons for transgenic rescue lines; three biologically independent experiments. Horizontal and vertical bars: Mean and SEM. Functional *osm-3* sequences were expressed in ADL and ADF under the *srh-220* and *srh-142* promoters, respectively. Data from two independent transgenic lines (l1, l2) are shown. ***: different from wildtype at P<0.001; #, ##, and ###: different from *osm-3* at P<0.05, 0.01, and 0.001, respectively; n.s.: not significant (Kruskal-Wallis test with Dunn’s correction for multiple comparisons). **e)** Percentage of neuron pairs exhibiting contacts between ADF and ADL cilia in animals of the indicated genotypes. Numbers indicate the number of neuron pairs examined; three biologically independent experiments. Data from two independent transgenic lines (l1, l2) are shown. ***: different from wildtype at P<0.001; #: different from *osm-3* at P<0.05 (Fisher’s exact tests with Holm’s correction for multiple comparisons). Errors are SEM. **f)** Extent of contacts between ADF and ADL cilia normalized for cilia length (see Methods). Each circle is the measurement from a single pair of neurons. n>23 neuron pairs; three biologically independent experiments. Data from two independent transgenic lines (l1, l2) are shown. Horizontal and vertical bars: Mean and SEM. ***: different from wildtype at P<0.001; n.s.: not significant (Kruskal-Wallis test with Dunn’s correction for multiple comparisons). *Alt text*: Images and quantification of ADL and ADF cilia length and contacts when their distal segments elongate in the absence of distal segments of other cilia in the amphid channel.

Since contacts are largely mediated by distal ciliary segments, we next tested whether distal segments of ADF and ADL cilia would be able to elongate and establish contact in *osm-3* mutants upon restoration of *osm-3* expression only in these neurons. In *osm-3(null)* mutants, distal segments of all channel cilia including those of ADF and ADL were truncated resulting in a complete loss of interciliary contacts (Fig. 2b-f) (Perkins et al. 1986; Signor et al. 1999). In contrast, loss of *kap-1* alone resulted in a slight shortening of the cilia of both neurons, with these neurons retaining partial interciliary contacts (Supplementary Fig. 1f-h). Expression of functional *osm-3* sequences in ADL and ADF alone was sufficient to partly but significantly elongate the cilia of both neurons in *osm-3(null)* mutants with weaker effects in ADF (Fig. 2b-d). We note, however, that these cilia often exhibited altered morphologies upon elongation (collectively referred to as ‘disorganized’ here) (Supplementary Fig. 2). These included cilia that exhibited distinct elongation trajectories, the two cilia of either ADL or ADF failing to separate (‘collapsed’ cilia), or elongation of only one cilium (Supplementary Fig. 2). Nonetheless, of the subset of ADF and ADL cilia whose length phenotype was rescued, a significant fraction established contacts of varying lengths including disorganized cilia (Fig. 2b,e,f). These results indicate that extension of the distal ciliary segments of ADL and to a lesser extent ADF, and establishment of interciliary contacts, can occur in the absence of the distal segments of other channel cilia, and that cilia exhibiting altered morphologies can retain the ability to form contacts.

### Cilia of monociliated neurons maintain their contact patterns in the absence of other amphid channel cilia

We next determined whether the cilia of monociliated neurons in the amphid channel are able to elongate and establish contacts in the absence of other channel cilia. Unlike in ADL and ADF, expression of functional *osm-3* sequences in the monociliated ASI or ASE neurons of *kap-1; osm-3* double mutants resulted in elongation of their single cilia (Supplementary Fig. 3a-d). While the cilia of ASH and ASI neurons do not contact each other, cilia of both these neurons are in direct contact with the cilia of ASE in the distal channel (Fig. 1b). Restoration of *osm-3* function to ASH/ASI and ASE neurons together in the *kap-1; osm-3* double mutant background was sufficient to not only rescue their cilia length defect, but to also fully rescue their interciliary contact length (Fig. 3a,c,d,e). In contrast, while expression of functional *osm-3* in ASH and ASI together also elongated both cilia, these cilia formed significantly shorter contacts largely at their distal ends similar to wild-type cilia (Fig. 3b,c-e). We were unable to fully resolve ASH-ASI cilia distance in the distal amphid channel by light microscopy, likely causing these cilia to appear to contact each other over a short distance in the narrow distal segments in contrast to observations using TEM. Restoration of *osm-3* function in ASH/ASI and ASE did not elongate ADL and ADF cilia, and conversely, *osm-3* expression in ADL and ADF had no effect on ASH and ASE cilia length, confirming the cell-specificity of *osm-3* expression (Supplementary Fig. 3e-h). Thus, elongation of the single rod-like cilia of a subset of amphid sensory neurons can occur in the absence of any other cilia in the channel. Moreover, these observations indicate that the formation of interciliary contact patterns between monociliated neurons do not require other adjacent cilia, and are unlikely to occur simply due to physical proximity.

**Fig. 3.**
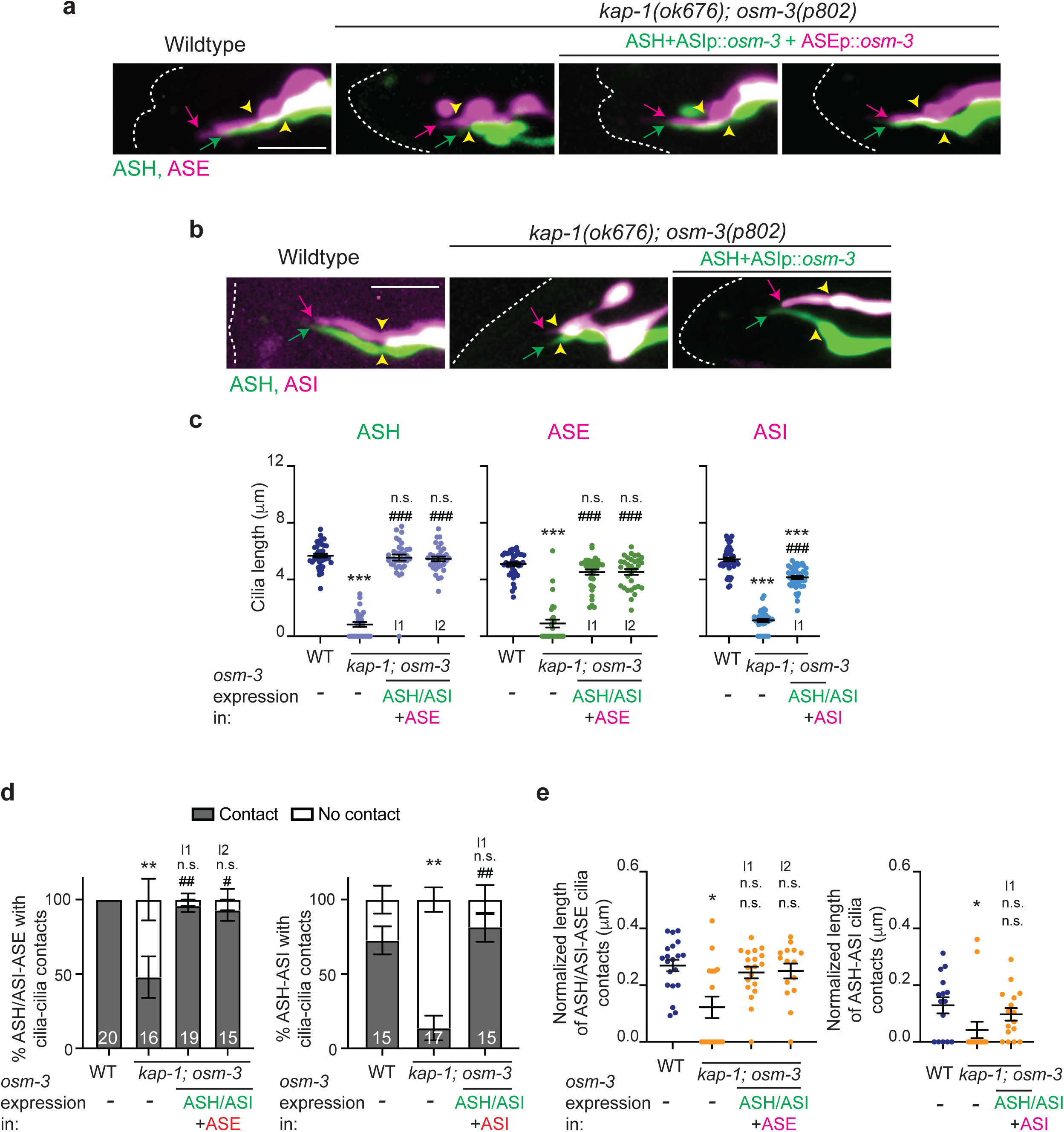
The pattern of interciliary contacts between monociliated neurons is maintained in the absence of other amphid channel cilia. **a,b)** Representative maximum intensity projection images of ASH/ASI and ASE (a), and ASH and ASI (b) cilia in the indicated genetic backgrounds. Dashed white lines: worm nose/head; arrows: cilia tip; yellow arrowheads: cilia base. Anterior at left; oriented to visualize both amphid channels (see Methods). Scale bar (for all images): 5 μm. **c)** Length of indicated cilia in animals with the indicated genetic backgrounds. Each circle is the length of a single cilium of each neuron. n>28 neurons for ASH, ASE, and ASI, n>39 neurons for transgenic rescue lines; three biologically independent experiments. Horizontal and vertical bars: Mean and SEM. Functional *osm-3* sequences were expressed in ASH/ASI, ASI and ASE under the *sra-6*, *srg-47,* and *flp-6* promoters, respectively. l1, l2 indicate independent transgenic lines. ***: different from wildtype at P<0.001; ###: different from mutant at P<0.001; n.s.: not significant (Kruskal-Wallis test with Dunn’s correction for multiple comparisons). **d)** Percentage of neuron pairs exhibiting contacts between ASH/ASI and ASE (left) and ASH and ASI (right) cilia in animals of the indicated genotypes. Numbers indicate the number of neuron pairs examined; three biologically independent experiments. l1, l2 indicate independent transgenic lines. **: different from wildtype at P<0.01; # and ##: different from mutant at P<0.05 and 0.01, respectively; n.s.: not significant (Fisher’s exact tests with Holm’s correction for multiple comparisons). Errors are SEM. **e)** Extent of contacts between ASH/ASI and ASE cilia, and ASH and ASI cilia normalized for length. Each circle is the measurement from a single pair of neurons. n>15 neuron pairs; three biologically independent experiments. l1, l2 indicate independent transgenic lines. Horizontal and vertical bars: Mean and SEM. *: different from wildtype at P<0.05; n.s.: not significant (Kruskal-Wallis test with Dunn’s correction for multiple comparisons). *Alt text*: Images and quantification of ASH, ASI, and ASE cilia length and contacts when they elongate in the absence of other cilia in the amphid channel.

### Dendrite order in the amphid neuron bundle does not dictate cilia contacts in the amphid channel

The dendrites of sensory neurons are organized in a stereotyped order in the amphid bundle, and these dendrites enter the amphid channel at specific locations (Doroquez et al. 2014; Ward et al. 1975; Yip and Heiman 2018). Although the absence of cilia does not alter dendrite order (Yip and Heiman 2018), we considered the possibility that the order and/or entry points of dendrites into the amphid channel may dictate cilia order and contacts. To address this notion, we examined mutants in which the relative dendrite order in the amphid bundle is altered.

Mutations in the *sax-7* L1 cell adhesion molecule gene disrupt relative dendrite order in the amphid neuron bundle in adult animals (Yip and Heiman 2018), and this protein is restricted to sensory dendrites (Lillis et al. 2022; Low et al. 2019). However, despite this disorganization, the distance between the bases of ADL and ADF cilia, as well as the location of the cilia bases in the amphid channel relative to the nose, were unaltered or affected to a relatively minor extent in *sax-7(eq1)* mutants (Fig. 4a-c). In addition, the percentage of ADL and ADF cilia exhibiting contacts and the extent of contact were unaffected in *sax-7* mutants (Fig. 4a, d-e). These observations suggest that dendrite order does not directly regulate cilia order in the amphid channel.

**Fig. 4.**
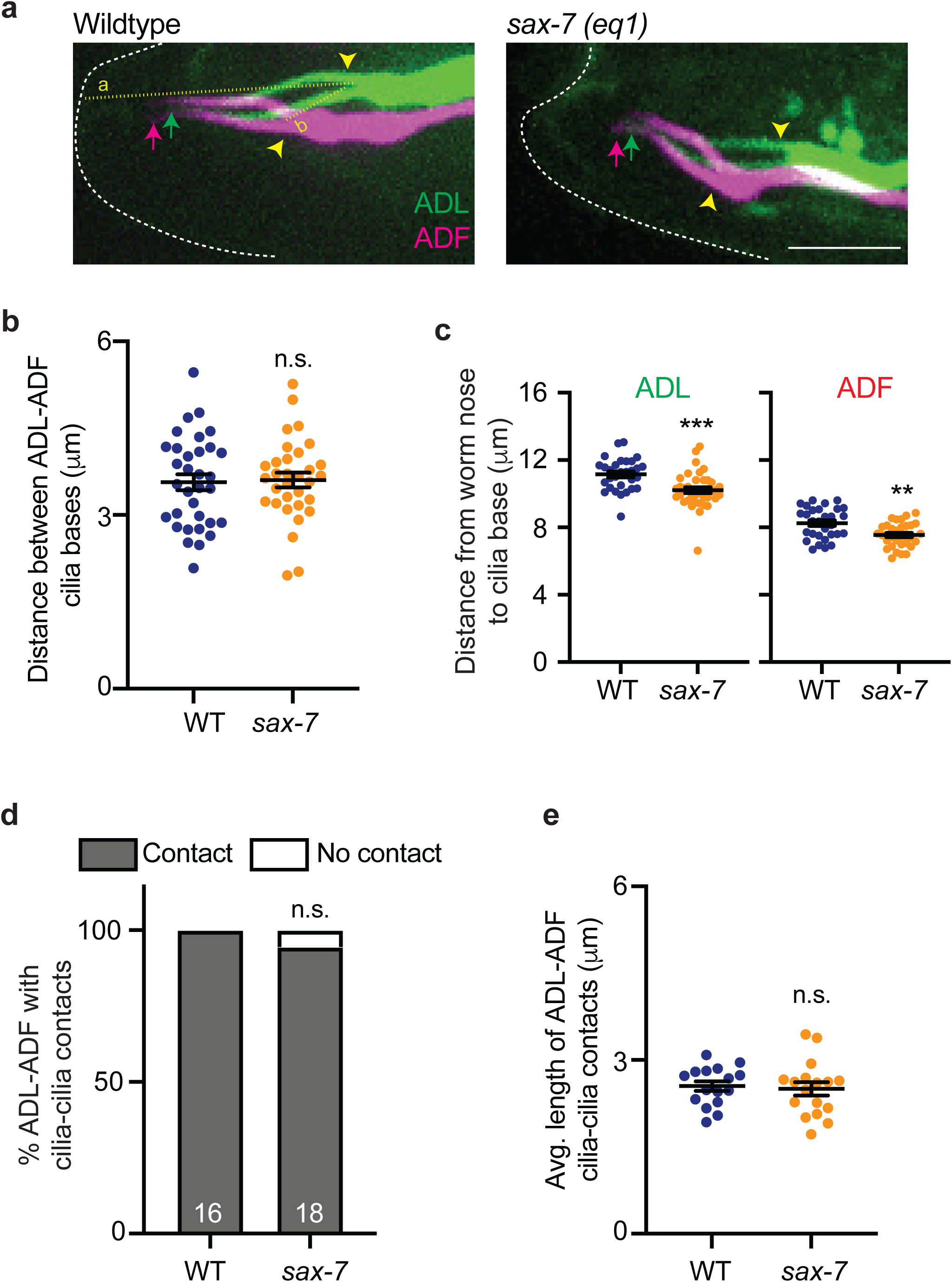
Disruption of dendrite order in the amphid neuron bundle does not alter contacts between ADL and ADF cilia. **a)** Representative maximum intensity projection images of ADL and ADF cilia in wildtype (WT) and *sax-7(eq1)* animals. Dashed white line: worm nose; arrows: cilia tip; yellow arrowheads: cilia base. Yellow dotted lines: a - distance of the cilia base from the worm nose (only shown for ADL), b - distance between ADL and ADF cilia bases. Anterior at left. Scale bar (for both images): 5 μm. **b,c)** Distance between bases of ADL and ADF cilia (b) and distance of ADL (c, left) and ADF (c, right) cilia bases from the worm nose in the indicated genetic backgrounds. Each circle is the measurement from a single pair of neurons (b) or each neuron (c). n >31 neuron pairs or neurons; two biologically independent experiments. Horizontal and vertical bars: Mean and SEM. ** and ***: different from corresponding wildtype at P<0.01 and 0.001, respectively; n.s.: not significant (Mann-Whitney test). **d,e)** Percentage of ADL and ADF neuron pairs exhibiting contacts between cilia (d) and the extent of contacts (e). Numbers in d indicate the number of neuron pairs examined. Each circle in **e** is the measurement from a single pair of neurons. n >16 neuron pairs; two biologically independent experiments. Horizontal and vertical bars: Mean and SEM. n.s.: not significant (d: Fisher’s exact test, e: Mann-Whitney test). *Alt text:* Images and quantification showing that disruption of dendrite order does not alter cilia order in the amphid channel.

### Mutations in a subset of genes implicated in the regulation of ciliary membrane composition and trafficking alter interciliary contacts and/or cilia organization

Combinatorial expression of cell adhesion molecules mediates fasciculation, target choice and wiring in the nervous system across species (eg. (Gerrow and El-Husseini 2006; Meltzer and Schuldiner 2022; Missaire and Hindges 2015; Schmucker 2007; Shapiro et al. 2007; Verpoort and de Wit 2024). In *Chlamydomonas*, flagellar agglutinins have been implicated in flagellar adhesion during mating (Adair et al. 1983), and in *C. elegans*, changes in membrane phospholipid composition have been suggested to alter ciliary contacts (Turan et al. 2025). We thus tested whether mutations in genes implicated in regulating ciliary membrane composition and ciliary protein trafficking affect interciliary contacts.

Pharmacological inhibition of the maturation of N-linked glycoproteins was shown to disrupt interciliary contacts among cultured MDCK cells (Ott et al. 2012), and altered N-glycosylation of flagellar proteins decreases flagellar adhesion in *Chlamydomonas* (Xu et al. 2020). ADL and ADF cilia lengths were weakly but significantly shortened in animals mutant for the conserved *aman-2* Golgi alpha-mannosidase II enzyme required for N-glycosylation (Rahman et al. 2022) (Fig. 5a,b), but the distance between the bases of ADF and ADL cilia, the location of the cilia bases relative to the nose, and the overall morphology of cilia were largely unaltered (Fig. 5c-e). We found that while there were also no defects in the percentage of cilia that contacted each other in *aman-2* mutants, when normalized for cilia length, these animals exhibited significant defects in the extent of interciliary contacts between ADF and ADL (Fig. 5f,g).

**Fig. 5.**
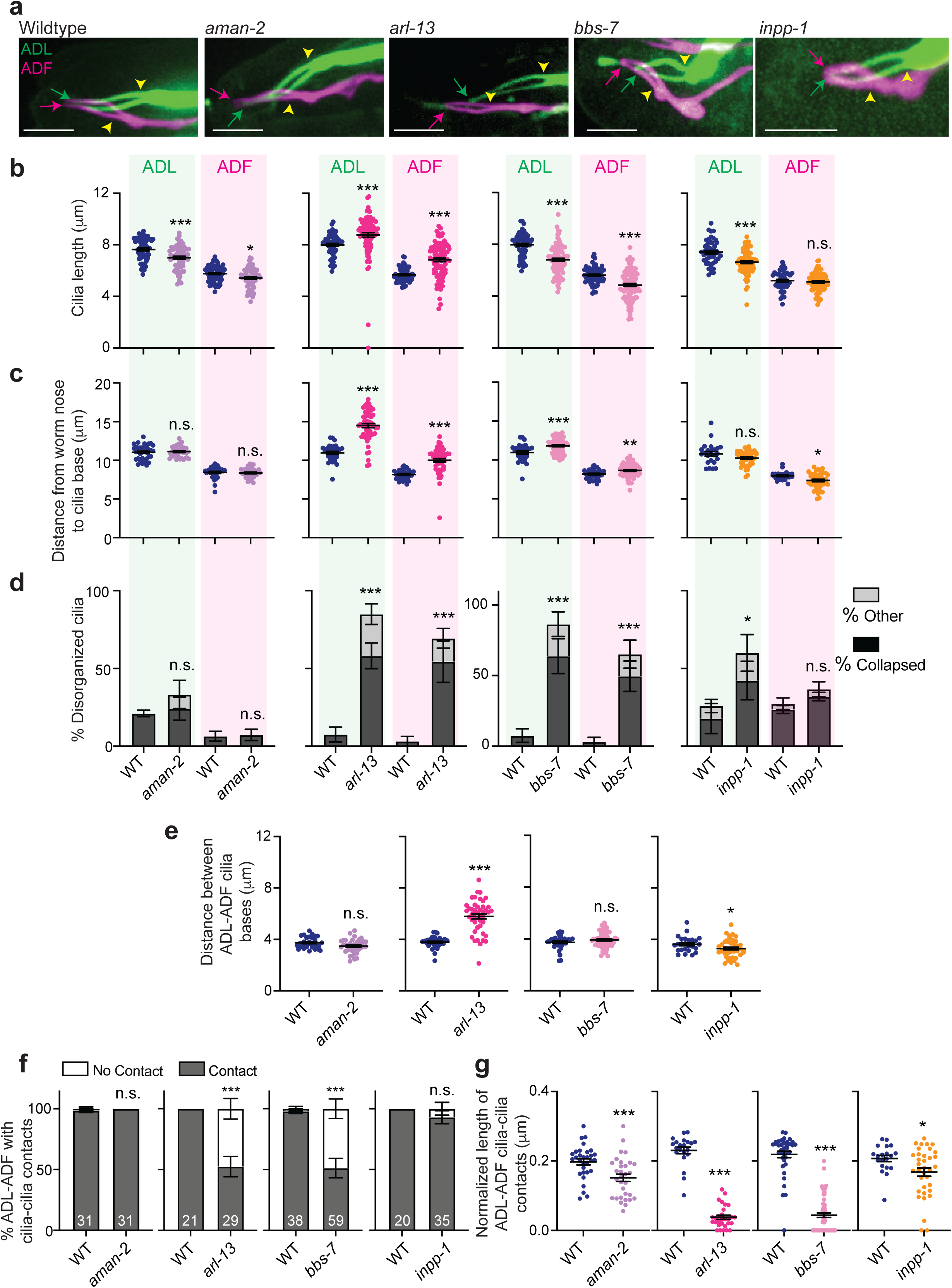
Mutations in genes required for ciliary membrane phospholipid composition or protein trafficking alter cilia organization and/or interciliary contacts. **a)** Representative maximum intensity projection images of ADF and ADL cilia in the indicated genetic backgrounds. Dashed white line: worm nose; arrows: cilia tip; yellow arrowheads: cilia base. Anterior at left. Scale bars: 5 μm. **b,c)** Cilia lengths (b) and distance of the cilia base from the nose (c) in ADL and ADF in the indicated genetic backgrounds. Each circle is the length of a single cilium (b) or distance from the nose (c) for a single neuron. n>49 (b), n>23 (c); three biologically independent experiments. **d)** Percentage of neurons exhibiting altered cilia morphology. ‘Collapsed’ refers to ADL or ADF cilia that fail to separate, ‘other’ refers to altered cilia morphologies including disrupted cilia trajectory and extension of a single cilium (see Supplementary Fig. 2). n>20 neurons; three biologically independent experiments. **e)** Distance between the ADL and ADF cilia bases in the indicated genetic backgrounds. Each dot is the quantification from a single neuron pair. n>25 neuron pairs; three biologically independent experiments. **f)** Percentage of ADL and ADF neurons exhibiting contacts between cilia in animals of the indicated genotypes. Numbers indicate the number of neuron pairs examined; three biologically independent experiments. **g)** Extent of contacts between ADF and ADL cilia normalized for cilia length (see Methods). Each circle is the measurement from a single pair of neurons. n>20 neuron pairs; three biologically independent experiments. Alleles used are indicated in Supplementary Table 1. Horizontal and vertical bars: Mean and SEM. *, ** and ***: different from corresponding wildtype at P<0.05, 0.01 or 0.001, respectively (b,c,e,g: Mann-Whitney test; d,f: Fisher’s exact test). n.s. - not significant. *Alt text*: Images and quantification showing that mutations in genes regulating cilia protein trafficking and membrane composition have differential effects on cilia morphology and interciliary contacts.

Transport of transmembrane and membrane-associated proteins in cilia is mediated in part via the BBSome complex and the ARL13 small GTPase (Blacque et al. 2004; Cevik et al. 2010; Fisher et al. 2020; Hor and Goh 2019; Li et al. 2012; Nachury 2018; Wingfield et al. 2018). *bbs* and *arl-13* mutants have also previously been implicated in cilia organization and elongation in *C. elegans* (Blacque et al. 2004; Cevik et al. 2010; Li et al. 2012; Turan et al. 2025). Moreover, ARL13 regulates ciliary membrane phospholipid composition via regulating ciliary localization of the INPP5E inositol polyphosphate 5-phosphatase (DiTirro et al. 2019; Humbert et al. 2012; Qiu et al. 2021; Turan et al. 2025). ADF and ADL cilia lengths were significantly longer in *arl-13* but shorter in animals mutant for the *bbs-7* BBSome complex component or the *inpp-1* INPP5E gene (Fig. 5a,b). The positions of the cilia bases were shifted distally in both *arl-13* and *bbs-7* mutants with weaker effects in *inpp-1* animals, although the distance between the ADL and ADF cilia bases was markedly increased only in *arl-13* mutants (Fig. 5c,e). Both *arl-13* and *bbs-7* but not *inpp-1* mutants exhibited strong defects in cilia organization with a large percentage of neurons containing cilia that failed to separate (‘collapsed’), as well as cilia that followed different trajectories (Fig. 5a,d). Correspondingly, both the fraction of contacting cilia as well as the extent of interciliary contacts were significantly reduced in *arl-13* and *bbs-7* mutants (Fig. 5f,g). We also observed defects in the extent of interciliary contact in *inpp-1* mutants although the percentage of cilia contacting each other was unaltered in this mutant background (Fig. 5f,g). We infer that while interciliary contact defects could arise as a consequence of, or be causal to, defects in cilia organization and morphology in *arl-13* and *bbs-7* mutants, the decreased interciliary contact length in *aman-2* and *inpp-1* mutants in the absence of significant defects in cilia organization support the hypothesis that interciliary contacts are mediated by specific interactions possibly via cilia-localized proteins.

### The BUG-1 secreted multidomain protein alters both cilia organization and interciliary contacts

We recently showed that the BUG-1 protein is completely required for attachment of the cilia of the BAG and URX sensory neurons to the IL socket glial cell (Wexler *et al*. 2026). *bug-1* encodes a secreted large protein containing CASH carbohydrate-binding, multiple EGF, and a Cadherin-like domains (Wexler *et al*. 2026). In addition to being expressed in BAG and URX, *bug-1* is also expressed in a subset of amphid channel sensory neurons and appears to be localized to cilia (Taylor et al. 2021). We tested the possibility that this molecule also mediates interciliary contacts in the amphid channel.

The previously described *bug-1(hmn404)* allele is a nonsense mutation in the 3^rd^ exon (Wexler *et al*. 2026). Cilia lengths of ADF and ADL were significantly elongated and the positions of the cilia bases were shifted distally in *bug-1(hmn404)* mutants (Fig. 6a-c, Supplementary Fig. 4a). The distance between ADF and ADL cilia bases was also more variable in *bug-1* mutants (Fig. 6e). Cilia organization was affected in the absence of *bug-1* with cilia of either or both ADF and ADL exhibiting defects in their morphologies in >90% of animals in this mutant background (Fig. 6d, Supplementary Fig. 4a). These defects included loss of one or both cilia, collapsed cilia, as well as cilia with markedly distinct trajectories (Fig. 6a,d, Supplementary Fig. 4a). Although morphologies of both neuronal cilia were typically altered in each amphid channel in *bug-1* mutants, we identified 6 ADL and 3 ADF cilia that exhibited altered morphology without effects on the corresponding neuronal cilia in the same channel implying that ciliary defects in these neurons may be regulated independently. A subset of cilia pairs in *bug-1* mutants failed to contact each other likely due to mispositioning in the amphid channel or cilia disorganization (Fig. 6f). Of the cilia pairs that established interciliary contact, the extent of contact was highly variable and on average shorter in *bug-1* mutants (Fig. 6g, Supplementary Fig. 4b). However, we noted that a subset of cilia exhibited significantly aberrant cilia morphology and trajectories in *bug-1* animals but still exhibited interciliary contacts (Figure 6a,g, Supplementary Fig. 4a). Thus, in contrast to URX and BAG cilia that are fully dependent on BUG-1 for their attachment to glia (Wexler *et al*. 2026), ADF and ADL cilia remain competent to form contacts in the absence of BUG-1 in some individuals. We suggest that BUG-1 is required for ADL and ADF cilia morphology and, directly or indirectly, also promotes contacts between ADL and ADF cilia.

**Fig. 6.**
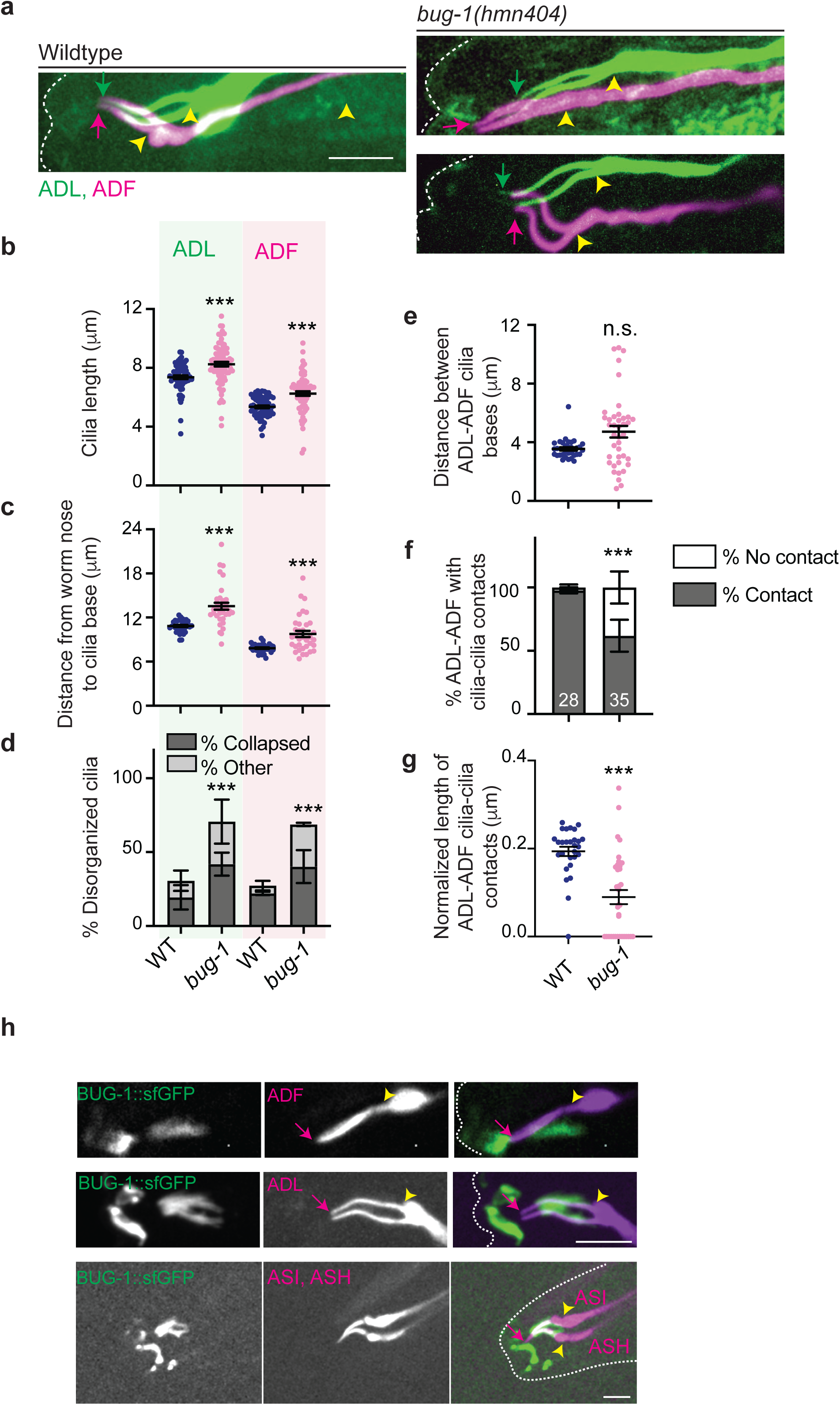
The BUG-1 multidomain protein localizes to the middle ciliary segments of a subset of amphid neurons and regulates cilia organization and interciliary contacts. **a)** Representative maximum intensity projection images of ADF and ADL cilia in wildtype and *bug-1* mutants. Dashed white line: worm nose; arrows: cilia tip; yellow arrowheads: cilia base. Anterior at left. Scale bar (for all images): 5 μm. Additional images of the ADF/ADL cilia phenotype in *bug-1* mutants are shown in Supplementary Fig. 4a. **b,c)** Cilia lengths (b) and distance of the cilia base from the nose (c) in ADL and ADF in wildtype and *bug-1* mutants. Each circle is the length of a single cilium (b) or distance from the nose (c) for each neuron. n>61 neurons (b), n>33 neurons (c); three biologically independent experiments. **d)** Percentage of neurons exhibiting altered cilia morphology. ‘Collapsed’ refers to ADL or ADF cilia that fail to separate, ‘other’ refers to altered cilia morphologies including disrupted cilia trajectory and extension of a single cilium. n>40 neurons; three independent experiments. Also see Supplementary Fig. 4a. **e)** Distance between the ADL and ADF cilia bases in the indicated genetic backgrounds. Each dot is the quantification from a single neuron pair. n>34 neuron pairs; three biologically independent experiments. **f)** Percentage of ADF and ADL neurons exhibiting contacts between cilia in animals of the indicated genotypes. Numbers indicate the number of neuron pairs examined; three biologically independent experiments. **g)** Extent of contacts between ADF and ADL cilia normalized for cilia length (see Methods). Each circle is the measurement from a single pair of neurons. n>28 neuron pairs; three biologically independent experiments. **h)** Representative maximum intensity projection images showing localization of an endogenously tagged BUG-1 fusion protein (Wexler *et al*. 2026) in the distal segments of ADL, ASH and ASI but not ADF cilia. ADL, ADF and ASH/ASI cilia were marked via expression of fluorescent reporters driven under the *srh-220, srh-142,* and *sra-6* promoters, respectively. Dashed white lines: worm nose/head; yellow arrowheads: cilia base; arrows: cilia tip. Anterior at left. Scale bar (for all images): 5 μm. Horizontal and vertical bars: Mean and SEM. ***: different from corresponding wildtype at P<0.001 (b,c,e,g: Mann-Whitney test; d,f: Fisher’s exact test). n.s. – not significant. *Alt text*: Images and quantification showing that mutations in the *bug-1*-encoded secreted protein alters cilia morphology and interciliary contacts.

Transcriptomics data from adult hermaphrodites indicate that among the amphid channel neurons, *bug-1* is expressed highly in ASH, ADL, and ASK with lower expression in ADF, ASI, and ASJ (Taylor et al. 2021) (www.cengen.org). Consistent with transcriptomics data, we observed that an endogenous reporter-tagged BUG-1 protein (Wexler *et al*. 2026) was associated with the middle ciliary segments of ADL, ASH and ASI, although we were unable to confirm association with ASJ and ASK (Fig. 6h). We did not detect association with ADF but are unable to exclude the possibility of low expression in this neuron type (Fig. 6h) (Taylor et al. 2021). In all expressing amphid sensory neurons, the BUG-1 fusion protein was present only around the middle ciliary segments (Fig. 6h). The absence of BUG-1 in the distal ciliary segments suggests that BUG-1 may regulate contacts between distal ciliary segments via indirect mechanisms.

## DISCUSSION

Here we show that the distal segments of a subset of sensory neuron cilia in *C. elegans* exhibit stereotyped organization and interciliary contacts in the amphid channel. Contact patterns between cilia pairs appear to be maintained in the absence of neighboring cilia, as well as in a subset of mutants that alter cilia trajectory, and are disrupted in mutants that are predicted to alter ciliary membrane composition and proteins, suggesting that these contacts may be established via directed cilia-cilia interactions. These interciliary connections may stabilize the cilia bundle architecture, and/or enable localized intercellular signaling.

During development, extension of a pioneer axon can form the scaffold for the extension of follower axons, thereby enabling the establishment of an ordered neuronal tract (Bate 1976; Ho and Goodman 1982). Channel sensory neurons are born between ∼300 and 450 minutes post fertilization with ASH, ADF, and ADL being the earliest and ASI the latest born neuron type (Sulston et al. 1983). The dendritic tips of these neurons are anchored at the nose, and dendrites elongate via retrograde extension, as the soma move posteriorly to their final positions (Fan et al. 2019; Heiman and Shaham 2009). Channel cilia appear to extend following dendritic extension in the embryo although the order of cilia growth in the channel during development is unknown (Nechipurenko et al. 2017; Serwas et al. 2017). Our observations argue against a model of a ‘pioneer cilium’ since we find that the cilia of multiple monociliated neurons are able to extend normally in the absence of other cilia in the channel in the adult animal. The exceptions are the biciliated ADL and ADF neurons whose cilia fail to extend in *osm-3; kap-1* double mutants upon cell-specific expression of *osm-3*. However, these cilia are able to extend when *osm-3* function is restored to all cilia in an *kap-1; osm-3(ts)* background suggesting that OSM-3-mediated elongation of these cilia requires the presence of one or more channel cilia as a scaffold. Loss of cilia upregulates glial secretion of extracellular matrix proteins into the amphid channel (Varandas et al. 2025; Wallace et al. 2016); it is possible that altered channel matrix composition inhibits elongation of biciliated but not monociliated channel neurons in the absence of other channel cilia. However, extension of the distal ciliary segments of ADF and ADL can occur without other distal ciliary segments in the channel, indicating that extension of the distal cilia does not require a pre-existing ciliary scaffold.

Three lines of evidence suggest that the pattern of interciliary contacts in the amphid channel may be mediated via an actively regulated process. First, contact patterns between cilia pairs appear to be maintained even in the absence of other cilia in the channel. For instance, the two cilia each of ADF and ADL and the single cilia of ASH/ASI and ASE are able to establish significant interciliary contacts when *osm-3* function is restored cell-specifically in *osm-3* or *kap-1; osm-3* double mutants despite the absence of distal segments or entire cilia, respectively, of all other channel cilia. In contrast, ASH and ASI cilia form minimal contacts under these conditions, suggesting that interciliary contacts are not formed indiscriminately. Second, mutations in genes such as *aman-2* and *inpp-1* that are predicted to alter ciliary glycoprotein or membrane phospholipid composition result in defects in the extent of interciliary overlap without major effects on cilia organization, suggesting that contacts may be mediated by cilia-localized proteins. ARL-13 and BBS-7 regulate ciliary trafficking of membrane proteins as well as membrane phospholipid composition (Blacque et al. 2004; Cevik et al. 2010; Fisher et al. 2020; Humbert et al. 2012; Li et al. 2012; Nachury 2018; Turan et al. 2025; Wingfield et al. 2018). Mutations in these genes disrupt channel cilia length, morphology, and interciliary contacts, and we suggest that these proteins may also regulate the transport or localization of ciliary proteins required for cilia-cilia contact. Third, a subset of ADL and ADF cilia that exhibit markedly distinct trajectories and thus likely have distinct physical neighbors, are nevertheless able to establish contacts with their correct partners in *bug-1* mutants albeit at a reduced frequency and length. Taken together, we propose that cilia contacts are mediated via instructive mechanisms. In future, it will be important to identify proteins that mediate these interciliary interactions and determine whether as in tissue morphogenesis or synapse formation (Honig and Shapiro 2020; Sanes and Zipursky 2020), combinatorial cell-specific expression of ciliary cell adhesion molecules instruct specific interciliary contact patterns.

Proteomic studies have identified adhesion molecules present in cilia including in neuronal cilia, and agglutinins are recruited to the flagella of mating *Chlamydomonas* (Chang et al. 2025; Hunnicutt et al. 1990; Ishikawa et al. 2012; Kohli et al. 2017; Liu et al. 2007; Macarelli et al. 2025; May et al. 2021; Mick et al. 2015). These molecules need not necessarily be localized specifically to the distal ciliary segments of channel cilia, since the physical distance between cilia may preclude interciliary contacts in the proximal channel (Doroquez et al. 2014). Intriguingly, each of the two cilia of ADF or ADL forms a partly distinct set of contacts (Doroquez et al. 2014), suggesting that each of these cilia may potentially localize a distinct set of adhesion molecules. In the visual circuits of *Drosophila* and mammals, as well as in the amphid neuron dendritic bundle of *C. elegans*, loss of contact with their partner results in a non-random order of connectivity with alternative targets suggesting a hierarchy of preferred interactions (Wolterhoff and Hiesinger 2024; Xu et al. 2019; Yip and Heiman 2018; Yoo et al. 2023; Zhang et al. 2022). Whether a similar hierarchy involving both attraction and repulsion underlies the formation of the observed contact patterns in the amphid channel remains to be determined.

Interflagellar adhesion triggers an intracellular signaling cascade during *Chlamydomonas* mating (Pasquale and Goodenough 1987). Cilia not only receive information, but may also actively transmit signals via the production of extracellular vesicles (Luxmi and King 2022; Wang and Barr 2018). The presence of connexins in neuronal cilia also suggests that cilia may communicate with each other via gap junctions, or influence the functions of neighboring neurons via non-synaptic modes of communication (Bokil et al. 2001; Su et al. 2012; Wu et al. 2024). In ctenophores, interciliary bridges connect adjacent motile cilia in their comb plates to coordinate ciliary beating (Dentler 1981). It remains possible that interactions among channel cilia contributes to the maintenance of ciliary integrity and/or function. Important goals for the future will be to identify the molecular mechanisms mediating interciliary interactions and to assess whether disruption of primary cilia contacts affects sensory encoding and behaviors in *C. elegans* and other organisms.

## Data availability

Strains and plasmids are available upon request. All strains used in this work are listed in Supplementary Table 1. All plasmids used in this work are listed in Supplementary Table 2. Data underlying analyses presented in each Figure are available at 10.6084/m9.figshare.31825261.

## Acknowledgements

We are grateful to the *Caenorhabditis* Genetics Center (CGC) for strains, Andrew Stone and Berith Isaac in the Brandeis University Light Microscopy and Electron Microscopy Core Facilities for advice regarding image analyses, and Stephen Nurrish, Revanth Sudhireddy, and Jihye Yeon for assistance with the generation of strains. We thank the Sengupta lab for advice, and Alison Philbrook and Priya Dutta for comments on the manuscript.

## REFERENCES

Adair WS, Hwang C, Goodenough UW. 1983. Identification and visualization of the sexual agglutinin from the mating-type plus flagellar membrane of *Chlamydomonas*. Cell. 33:183–193.

Agi E, Kulkarni A, Hiesinger PR. 2020. Neuronal strategies for meeting the right partner during brain wiring. Curr Opin Neurobiol. 63:1–8.

Bate CM. 1976. Pioneer neurones in an insect embryo. Nature. 260:54–56.

Blacque OE, Reardon MJ, Li C, McCarthy J, Mahjoub MR, Ansley SJ, Badano JL, Mah AK, Beales PL, Davidson WS et al. 2004. Loss of *C. elegans bbs-7* and *bbs-8* protein function results in cilia defects and compromised intraflagellar transport. Genes Dev. 18:1630–1642.

Bokil H, Laaris N, Blinder K, Ennis M, Keller A. 2001. Ephaptic interactions in the mammalian olfactory system. J Neurosci. 21:RC173.

Cevik S, Hori Y, Kaplan OI, Kida K, Toivenon T, Foley-Fisher C, Cottell D, Katada T, Kontani K, Blacque OE. 2010. Joubert syndrome arl13b functions at ciliary membranes and stabilizes protein transport in *Caenorhabditis elegans*. J Cell Biol. 188:953–969.

Chang C-H, Trinh VN, Lokesh NR, Montecinos CK, Pownall ME, Kalocsay M, Nachury MV. 2025. *In situ* proteomics unveils specialized domains for extrasynaptic signaling on neuronal cilia. Sci Adv. 23: eaed5548.

Cook SJ, Kalinski CA, Hobert O. 2023. Neuronal contact predicts connectivity in the *C. elegans* brain. Curr Biol. 33:2315–2320 e2312.

Dentler WL. 1981. Microtubule-membrane interactions in ctenophore swimming plate cilia. Tissue Cell. 13:197–208.

DiTirro D, Philbrook A, Rubino K, Sengupta P. 2019. The *Caenorhabditis elegans* tubby homolog dynamically modulates olfactory cilia membrane morphogenesis and phospholipid composition. Elife. 8:e48789.

Doroquez DB, Berciu C, Anderson JR, Sengupta P, Nicastro D. 2014. A high-resolution morphological and ultrastructural map of anterior sensory cilia and glia in *C. elegans*. eLife. 3:e01948.

Falk N, Losl M, Schroder N, Giessl A. 2015. Specialized cilia in mammalian sensory systems. Cells. 4:500–519.

Fan L, Kovacevic I, Heiman MG, Bao Z. 2019. A multicellular rosette-mediated collective dendrite extension. Elife. 8:e38065.

Ferkey DM, Sengupta P, L’Etoile ND. 2021. Chemosensory signal transduction in *Caenorhabditis elegans*. Genetics. 217:iyab004.

Fisher S, Kuna D, Caspary T, Kahn RA, Sztul E. 2020. Arf family gtpases with links to cilia. Am J Physiol Cell Physiol. 319:C404–C418.

Gerrow K, El-Husseini A. 2006. Cell adhesion molecules at the synapse. Front Biosci. 11:2400–2419.

Goodman MB, Sengupta P. 2019. How *Caenorhabditis elegans* senses mechanical stress, temperature, and other physical stimuli. Genetics. 212:25–51.

Gumbiner BM. 1996. Cell adhesion: The molecular basis of tissue architecture and morphogenesis. Cell. 84:345–357.

Heiman MG, Shaham S. 2009. *dex-1* and *dyf-7* establish sensory dendrite length by anchoring dendritic tips during cell migration. Cell. 137:344–355.

Ho RK, Goodman CS. 1982. Peripheral pathways are pioneered by an array of central and peripheral neurones in grasshopper embryos. Nature. 297:404–406.

Hor CH, Goh EL. 2019. Small gtpases in hedgehog signalling: Emerging insights into the disease mechanisms of *rab23*-mediated and *arl13b*-mediated ciliopathies. Curr Opin Genet Dev. 56:61–68.

Humbert MC, Weihbrecht K, Searby CC, Li Y, Pope RM, Sheffield VC, Seo S. 2012. Arl13b, pde6d, and cep164 form a functional network for inpp5e ciliary targeting. Proc Natl Acad Sci USA. 109:19691–19696.

Hunnicutt GR, Kosfiszer MG, Snell WJ. 1990. Cell body and flagellar agglutinins in *Chlamydomonas reinhardtii:* The cell body plasma membrane is a reservoir for agglutinins whose migration to the flagella is regulated by a functional barrier. J Cell Biol. 111:1605–1616.

Ishikawa H, Thompson J, Yates JR, 3rd, Marshall WF. 2012. Proteomic analysis of mammalian primary cilia. Curr Biol. 22:414–419.

Jurisch-Yaksi N, Wachten D, Gopalakrishnan J. 2024. The neuronal cilium - a highly diverse and dynamic organelle involved in sensory detection and neuromodulation. Trends Neurosci. 47:383–394.

Kohli P, Hohne M, Jungst C, Bertsch S, Ebert LK, Schauss AC, Benzing T, Rinschen MM, Schermer B. 2017. The ciliary membrane-associated proteome reveals actin-binding proteins as key components of cilia. EMBO Rep. 18:1521–1535.

Kremer JR, Mastronarde DN, McIntosh JR. 1996. Computer visualization of three-dimensional image data using imod. J Struct Biol. 116:71–76.

Li Y, Zhang Q, Wei Q, Zhang Y, Ling K, Hu J. 2012. Sumoylation of the small gtpase arl-13 promotes ciliary targeting of sensory receptors. J Cell Biol. 199:589–598.

Lillis M, Zaccardi NJ, Heiman MG. 2022. Axon-dendrite and apical-basolateral sorting in a single neuron. Genetics. 221: iyac036.

Liu Q, Tan G, Levenkova N, Li T, Pugh EN, Jr., Rux JJ, Speicher DW, Pierce EA. 2007. The proteome of the mouse photoreceptor sensory cilium complex. Mol Cell Proteomics. 6:1299–1317.

Low IIC, Williams CR, Chong MK, McLachlan IG, Wierbowski BM, Kolotuev I, Heiman MG. 2019. Morphogenesis of neurons and glia within an epithelium. Development. 146: dev171124.

Luxmi R, King SM. 2022. Cilia-derived vesicles: An ancient route for intercellular communication. Semin Cell Dev Biol. 129:82–92.

Macarelli V, Sroka TJ, Jansen JN, Axelsson U, Mick DU, Merkle FT. 2025. Proximity proteomics of primary cilia in human hypothalamic neurons. bioRxiv. 10.1101/2025.05.11.653368

May EA, Kalocsay M, D’Auriac IG, Schuster PS, Gygi SP, Nachury MV, Mick DU. 2021. Time-resolved proteomics profiling of the ciliary hedgehog response. J Cell Biol. 220:e202007207.

Meltzer H, Schuldiner O. 2022. Spatiotemporal control of neuronal remodeling by cell adhesion molecules: Insights from *Drosophila*. Front Neurosci. 16:897706.

Mick DU, Rodrigues RB, Leib RD, Adams CM, Chien AS, Gygi SP, Nachury MV. 2015. Proteomics of primary cilia by proximity labeling. Dev Cell. 35:497–512.

Missaire M, Hindges R. 2015. The role of cell adhesion molecules in visual circuit formation: From neurite outgrowth to maps and synaptic specificity. Dev Neurobiol. 75:569–583.

Monfared RV, Abdelkarim S, Derdeyn P, Chen K, Wu H, Leong K, Chang T, Lee J, Versales S, Nauli SM et al. 2025. Cilia in the brain display region-dependent oscillations of length and orientation. PLoS Biol. 23:e3003197.

Muller A, Klena N, Pang S, Garcia LEG, Topcheva O, Aurrecoechea Duran S, Sulaymankhil D, Seliskar M, Mziaut H, Schoniger E et al. 2024. Structure, interaction and nervous connectivity of beta cell primary cilia. Nat Commun. 15:9168.

Nachury MV. 2018. The molecular machines that traffic signaling receptors into and out of cilia. Curr Opin Cell Biol. 51:124–131.

Nechipurenko IV, Berciu C, Sengupta P, Nicastro D. 2017. Centriolar remodeling underlies basal body maturation during ciliogenesis in *Caenorhabditis elegans*. Elife. 6:e25686.

Ott C, Elia N, Jeong SY, Insinna C, Sengupta P, Lippincott-Schwartz J. 2012. Primary cilia utilize glycoprotein-dependent adhesion mechanisms to stabilize long-lasting cilia-cilia contacts. Cilia. 1:3.

Ott CM, Torres R, Kuan TS, Kuan A, Buchanan J, Elabbady L, Seshamani S, Bodor AL, Collman F, Bock DD et al. 2024. Ultrastructural differences impact cilia shape and external exposure across cell classes in the visual cortex. Curr Biol. 34:2418–2433 e2414.

Packer AM, McConnell DJ, Fino E, Yuste R. 2013. Axo-dendritic overlap and laminar projection can explain interneuron connectivity to pyramidal cells. Cerebr cortex. 23:2790–2802.

Pasquale SM, Goodenough UW. 1987. Cyclic amp functions as a primary sexual signal in gametes of *chlamydomonas reinhardtii*. J Cell Biol. 10:2279–2292.

Perkins LA, Hedgecock EM, Thomson JN, Culotti JG. 1986. Mutant sensory cilia in the nematode *caenorhabditis elegans*. Dev Biol. 117:456–487.

Peters A, Feldman ML. 1976. The projection of the lateral geniculate nucleus to area 17 of the rat cerebral cortex. I. General description. J Neurocytol. 5:63–84.

Philbrook A, O’Donnell MP, Grunenkovaite L, Sengupta P. 2024. Cilia structure and intraflagellar transport differentially regulate sensory response dynamics within and between *C. elegans* chemosensory neurons. PLoS Biol. 22:e3002892.

Qiu H, Fujisawa S, Nozaki S, Katoh Y, Nakayama K. 2021. Interaction of inpp5e with arl13b is essential for its ciliary membrane retention but dispensable for its ciliary entry. Biol Open. 10:bio058843.

Rahman M, Ramirez-Suarez NJ, Diaz-Balzac CA, Bulow HE. 2022. Specific n-glycans regulate an extracellular adhesion complex during somatosensory dendrite patterning. EMBO Rep. 23:e54163.

Rees CL, Moradi K, Ascoli GA. 2017. Weighing the evidence in Peters’ rule: Does neuronal morphology predict connectivity? Trends Neurosci. 40:63–71.

Sanes JR, Zipursky SL. 2020. Synaptic specificity, recognition molecules, and assembly of neural circuits. Cell. 181:536–556.

Schmucker D. 2007. Molecular diversity of dscam: Recognition of molecular identity in neuronal wiring. Nat Rev Neurosci. 8:915–920.

Serwas D, Su TY, Roessler M, Wang S, Dammermann A. 2017. Centrioles initiate cilia assembly but are dispensable for maturation and maintenance in *C. elegans*. J Cell Biol. 216:1659–1671.

Shapiro L, Love J, Colman DR. 2007. Adhesion molecules in the nervous system: Structural insights into function and diversity. Annu Rev Neurosci. 30:451–474.

Sheu SH, Upadhyayula S, Dupuy V, Pang S, Deng F, Wan J, Walpita D, Pasolli HA, Houser J, Sanchez-Martinez S et al. 2022. A serotonergic axon-cilium synapse drives nuclear signaling to alter chromatin accessibility. Cell. 185:3390–3407 e3318.

Signor D, Wedaman KP, Rose LS, Scholey JM. 1999. Two heteromeric kinesin complexes in chemosensory neurons and sensory cilia of *Caenorhabditis elegans*. Mol Biol Cell. 10:345–360.

Singla V, Reiter JF. 2006. The primary cilium as the cell’s antenna: Signaling at a sensory organelle. Science. 313:629–633.

Snow JJ, Ou G, Gunnarson AL, Walker MR, Zhou HM, Brust-Mascher I, Scholey JM. 2004. Two anterograde intraflagellar transport motors cooperate to build sensory cilia on *C. elegans* neurons. Nat Cell Biol. 6:1109–1113.

Sperry RW. 1963. Chemoaffinity in the orderly growth of nerve fiber patterns and connections. Proc Natl Acad Sci USA. 50:703–710.

Steinberg MS, Takeichi M. 1994. Experimental specification of cell sorting, tissue spreading, and specific spatial patterning by quantitative differences in cadherin expression. Proc Natl Acad Sci USA. 91:206–209.

Su CY, Menuz K, Reisert J, Carlson JR. 2012. Non-synaptic inhibition between grouped neurons in an olfactory circuit. Nature. 492:66–71.

Sulston JE, Schierenberg E, White JG, Thomson JN. 1983. The embryonic cell lineage of the nematode *Caenorhabditis elegans*. Dev Biol. 100:64–119.

Taylor SR, Santpere G, Weinreb A, Barrett A, Reilly MB, Xu C, Varol E, Oikonomou P, Glenwinkel L, McWhirter R et al. 2021. Molecular topography of an entire nervous system. Cell. 184:4329–4347 e4323.

Togashi H, Sakisaka T, Takai Y. 2009. Cell adhesion molecules in the central nervous system. Cell Adh Migr. 3:29–35.

Townes PL, Holftreter J. 1955. Directed movements and selective adhesion of embryonic amphibian cells. J Exp Zool. 128:53–120.

Turan MG, Kantarci H, Cevik S, Kaplan OI. 2025. Arl13b regulates juxtaposed cilia-cilia elongation in bbsome dependent manner in *Caenorhabditis elegans*. iScience. 28:111791.

Varandas KC, Hodges BM, Lubeck L, Farinas A, Liang Y, Lu Y, Shaham S. 2025. Glia detect and transiently protect against dendrite substructure disruption in *C. elegans*. Nat Commun. 16:79.

Verpoort B, de Wit J. 2024. Cell adhesion molecule signaling at the synapse: Beyond the scaffold. Cold Spring Harb Perspect Biol. 16:a041501.

Wallace SW, Singhvi A, Liang Y, Lu Y, Shaham S. 2016. Pros-1/prospero is a major regulator of the glia-specific secretome controlling sensory-neuron shape and function in *C. elegans*. Cell reports. 15:550–562.

Wang J, Barr MM. 2018. Cell-cell communication via ciliary extracellular vesicles: Clues from model systems. Essays Biochem. 62:205–213.

Ward S, Thomson N, White JG, Brenner S. 1975. Electron microscopical reconstruction of the anterior sensory anatomy of the nematode *caenorhabditis elegans*. J Comp Neurol. 160:313–337.

Wingfield JL, Lechtreck KF, Lorentzen E. 2018. Trafficking of ciliary membrane proteins by the intraflagellar transport/bbsome machinery. Essays Biochem. 62:753–763.

Wolterhoff N, Hiesinger PR. 2024. Synaptic promiscuity in brain development. Curr Biol. 34:R102–R116.

Wu JY, Cho SJ, Descant K, Li PH, Shapson-Coe A, Januszewski M, Berger DR, Meyer C, Casingal C, Huda A et al. 2024. Mapping of neuronal and glial primary cilia contactome and connectome in the human cerebral cortex. Neuron. 112:41–55.

Xu C, Theisen E, Maloney R, Peng J, Santiago I, Yapp C, Werkhoven Z, Rumbaut E, Shum B, Tarnogorska D et al. 2019. Control of synaptic specificity by establishing a relative preference for synaptic partners. Neuron. 103:865–877.

Xu N, Oltmanns A, Zhao L, Girot A, Karimi M, Hoepfner L, Kelterborn S, Scholz M, Beissel J, Hegemann P et al. 2020. Altered n-glycan composition impacts flagella-mediated adhesion in *Chlamydomonas reinhardtii*. Elife. 9:e58805.

Yip ZC, Heiman MG. 2018. Ordered arrangement of dendrites within a *C. elegans* sensory nerve bundle. Elife. 7:e35825.

Yoo J, Dombrovski M, Mirshahidi P, Nern A, LoCascio SA, Zipursky SL, Kurmangaliyev YZ. 2023. Brain wiring determinants uncovered by integrating connectomes and transcriptomes. Curr Biol. 33:3998–4005 e3996.

Zhang C, Hellevik A, Takeuchi S, Wong RO. 2022. Hierarchical partner selection shapes rod-cone pathway specificity in the inner retina. iScience. 25:105032.

