## Supplemental Figures, Legends, Tables for "Stereotypical interciliary contacts in a *C. elegans* sense organ"

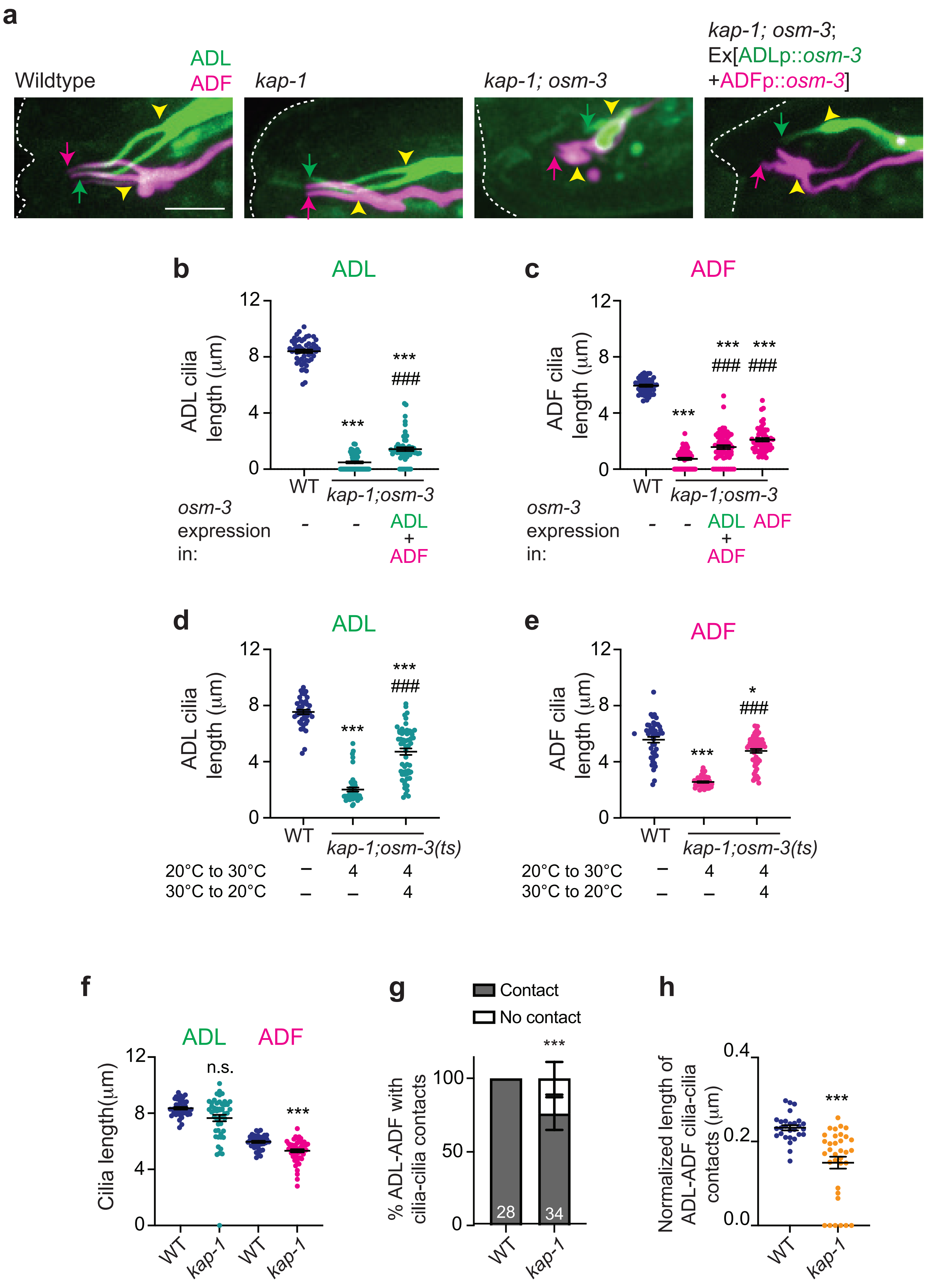
a
Wildtype
ADL
ADF
kap-1
kap-1; osm-3
kap-1; osm-3;
Ex[ADLp::osm-3
+ADFp::osm-3]
b
ADL
ADL cilia length (μm)
WT
kap-1;osm-3
osm-3 expression in:
ADL + ADF
c
ADF
ADF cilia length (μm)
WT
kap-1;osm-3
osm-3 expression in:
ADL + ADF
ADF
d
ADL
ADL cilia length (μm)
WT
kap-1;osm-3(ts)
20°C to 30°C
30°C to 20°C
e
ADF
ADF cilia length (μm)
WT
kap-1;osm-3(ts)
20°C to 30°C
30°C to 20°C
f
ADL
ADF
Cilia length(μm)
n.s.
WT
kap-1
g
Contact
No contact
% ADL-ADF with cilia-cilia contacts
28
34
WT
kap-1
h
Normalized length of ADL-ADF cilia-cilia contacts (μm)
WT
kap-1

**Supplementary Fig. 1. OSM-3-mediated elongation of ADL and ADF cilia in *kap-1; osm-3* double mutants requires other amphid channel cilia.**

**a)** Representative maximum intensity projection images of ADL and ADF cilia in animals of the indicated genotypes and under shown conditions. Alleles used were *kap-1(ok676)* and *osm-3(p802)*. Dashed white lines: worm nose; arrows: cilia tip; yellow arrowheads: cilia base. Anterior at left. Scale bar (for all images): 5 µm.

**b-f)** Length of ADL or ADF cilia in animals with the indicated genetic backgrounds and under the shown conditions. Each circle is the length of a single cilium of each neuron. n>24 neurons; two biologically independent experiments. Functional *osm-3* sequences were expressed in ADL and ADF under the *srh-220* and *srh-142* promoters, respectively in b and c. The *kap-1(ok676)* and *osm-3(oy156ts)* alleles were used in d and e. Horizontal and vertical bars: Mean and SEM. * and ***: different from corresponding wildtype at P<0.05 and 001, respectively; ###: different from mutant at P<0.001; n.s.: not significant (b,c: Kruskal-Wallis test with Dunn's correction for multiple comparisons; d: Mann-Whitney test).

**g,h)** Percentage of ADF and ADL neurons exhibiting contacts between cilia (g) and the extent of the contacts normalized for cilia length (h) in animals of the indicated genetic backgrounds and shown conditions. Numbers in g indicate the number of neuron pairs examined. Each circle in h is the measurement from a single pair of neurons. n>28 neuron pairs; three biologically independent experiments. Horizontal and vertical bars: Mean and SEM. ***: different from wildtype at P<0.001 (e: Fisher's exact test; f: Mann-Whitney test).

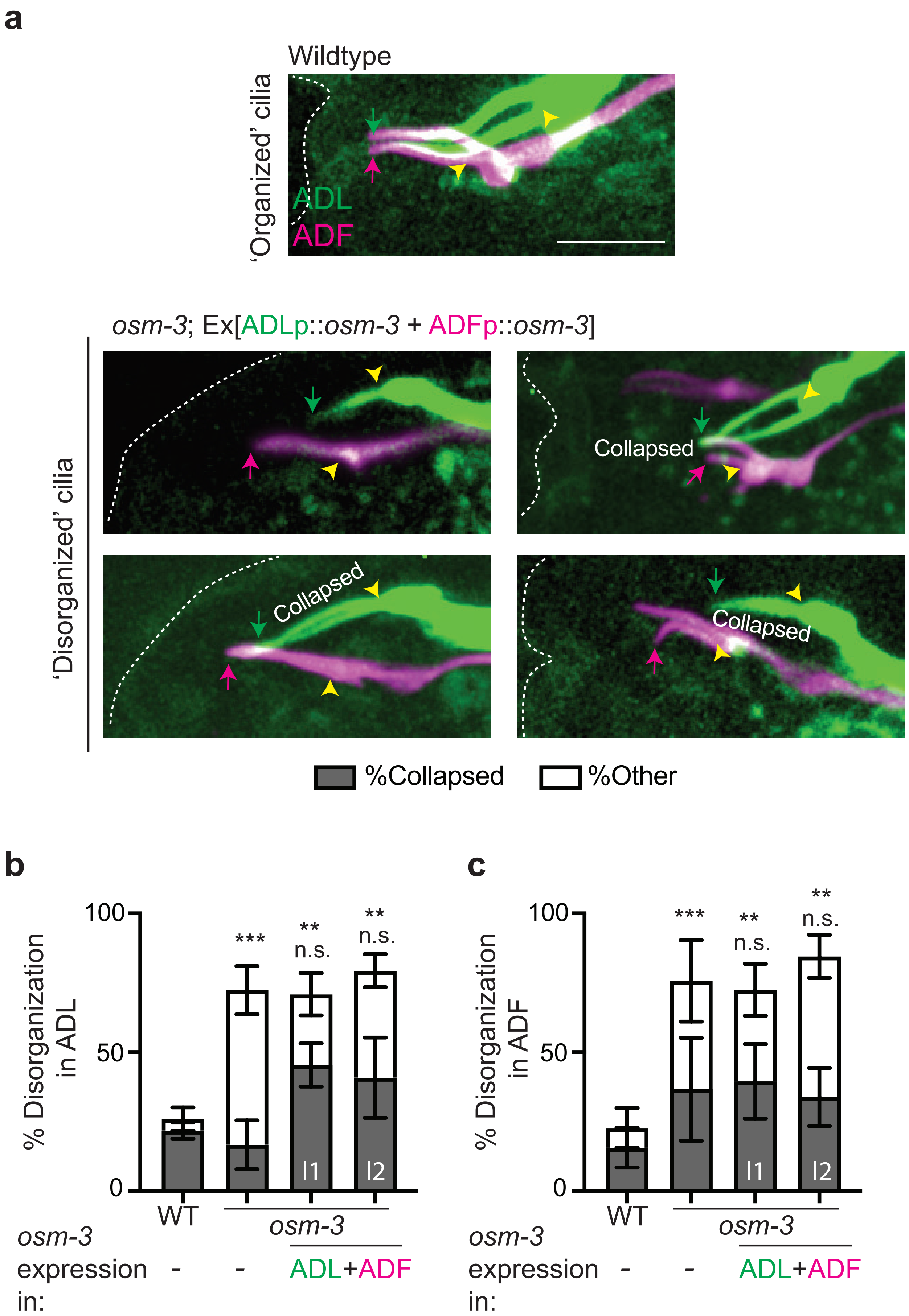
a
Wildtype
'Organized' cilia
ADL
ADF
osm-3; Ex[ADLp::osm-3 + ADFp::osm-3]
'Disorganized' cilia
Collapsed
Collapsed
Collapsed
%Collapsed
%Other
b
% Disorganization in ADL
100
50
0
***
**
n.s.
**
n.s.
l1
l2
WT
osm-3
osm-3 expression in:
-
-
ADL+ADF
c
% Disorganization in ADF
100
50
0
***
**
n.s.
**
n.s.
l1
l2
WT
osm-3
osm-3 expression in:
-
-
ADL+ADF

**Supplementary Fig. 2. Cilia morphologies of ADF and ADL are altered in the absence of the distal segments of other channel cilia.**

**a)** Representative maximum intensity projection images of ADL and ADF cilia showing the ‘organized’ phenotype in wildtype animals (top), and distinct ‘disorganized’ morphologies in *osm-3(p802)* mutants upon expression of functional *osm-3* sequences in ADL and ADF under the *srh-220* and *srh-142* promoters, respectively (bottom). ‘Disorganized’ includes cilia that exhibit distinct trajectories, elongation of a single cilium, and cilia of ADL or ADF that fail to separate (‘collapsed’). Dashed white line: worm nose; arrows: cilia tip; yellow arrowheads: cilia base. Anterior at left. Scale bar (for all images): 5 µm.

**b,c)** Percentage of ADL (b) and ADF (c) neurons in animals of the indicated genotypes exhibiting altered cilia morphology. n>23 neurons; three biologically independent experiments. Data from two independent transgenic lines (l1, l2) expressing functional *osm-3 s*equences in ADL and ADF are shown. Errors are SEM. **, and ***: different from corresponding wildtype at P<0.01 and 0.001, respectively; n.s.: not significant (Fisher’s exact tests with Holm’s correction for multiple comparisons).

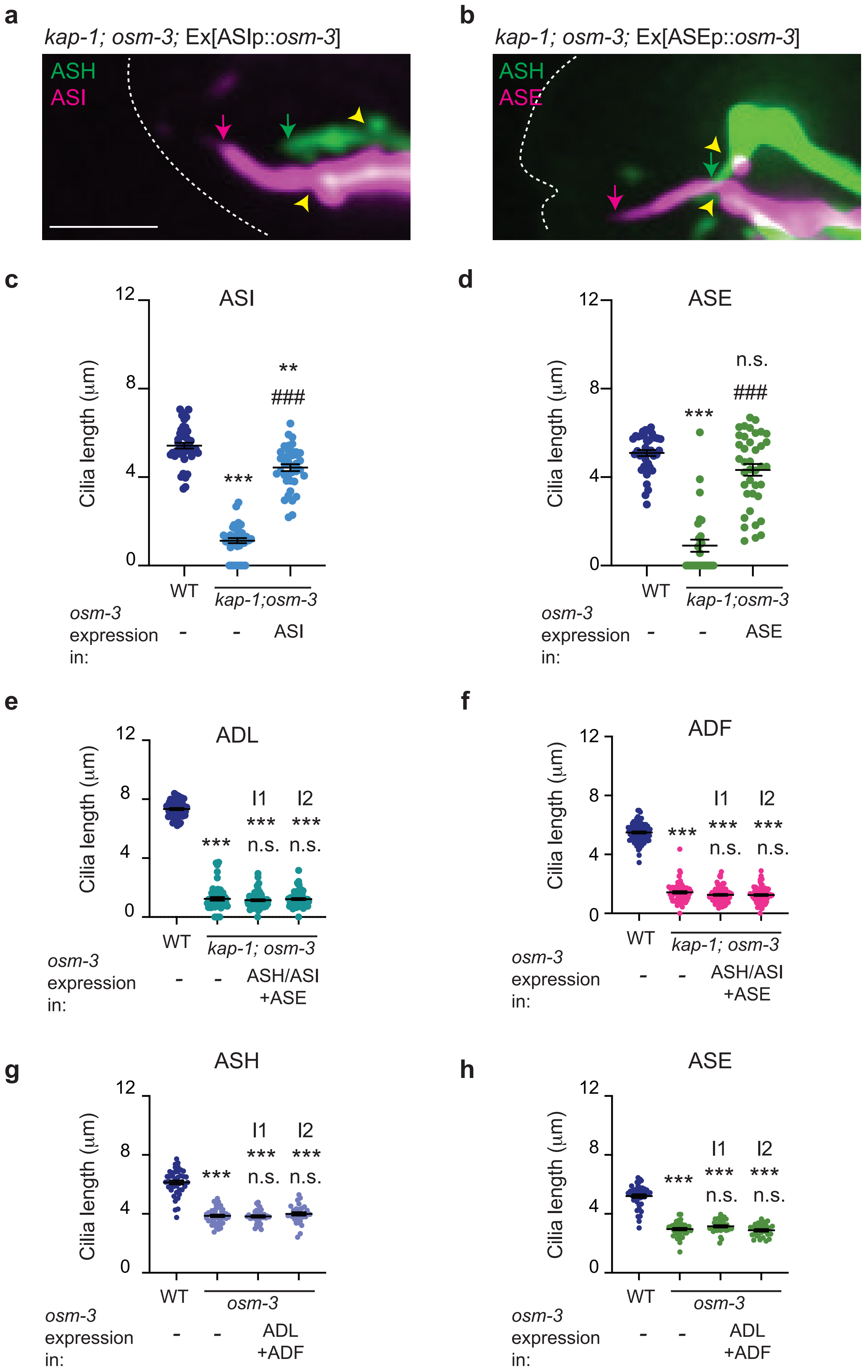

a
kap-1; osm-3; Ex[ASIp::osm-3]
ASH
ASI
b
kap-1; osm-3; Ex[ASEp::osm-3]
ASH
ASE
c
ASI
Cilia length (µm)
12
8
4
0
***
**
###
WT
kap-1;osm-3
osm-3 expression in:
-
-
ASI
d
ASE
Cilia length (µm)
12
8
4
0
***
n.s.
###
WT
kap-1;osm-3
osm-3 expression in:
-
-
ASE
e
ADL
Cilia length (µm)
12
8
4
0
***
I1
***
n.s.
I2
***
n.s.
WT
kap-1; osm-3
osm-3 expression in:
-
-
ASH/ASI +ASE
f
ADF
Cilia length (µm)
12
8
4
0
***
I1
***
n.s.
I2
***
n.s.
WT
kap-1; osm-3
osm-3 expression in:
-
-
ASH/ASI +ASE
g
ASH
Cilia length (µm)
12
8
4
0
***
I1
***
n.s.
I2
***
n.s.
WT
osm-3
osm-3 expression in:
-
-
ADL +ADF
h
ASE
Cilia length (µm)
12
8
4
0
***
I1
***
n.s.
I2
***
n.s.
WT
osm-3
osm-3 expression in:
-
-
ADL +ADF

**Supplementary Fig. 3. *osm-3* expression in the ASI and ASE monociliated neurons rescues their cilia length defect in *kap-1; osm-3* double mutants.**

**a,b)** Representative maximum intensity projection images of ASI (a) and ASE (b) cilia in *kap-1; osm-3* double mutants expressing functional *osm-3* in ASI and ASE, respectively. *osm-3* was expressed under the *srg-47* (ASI) and *flp-6* (ASE) promoters. The ASH neurons are also marked with a fluorescent reporter in these strains. Dashed white line: worm nose; arrows: cilia tip; yellow arrowheads: cilia base. Anterior at left. Scale bar (for both images): 5 μm.

**c-h)** Length of indicated cilia in animals with the shown genetic backgrounds. Each circle is the cilium length in a single neuron. n>27 neurons; two biologically independent experiments. Functional *osm-3* sequences were expressed in ASI, ASE, ASH/ASI, ADL and ADF under the *srg-47*, *flp-6*, *sra-6, srh-220*, and *srh-142* promoters, respectively. l1 and l2 indicate independent transgenic lines. Horizontal and vertical bars: Mean and SEM. ** and ***: different from corresponding wildtype at P<0.01 and 0.001; ###: different from mutant background at P<0.001; n.s.: not significant (Kruskal-Wallis test with Dunn's correction for multiple comparisons).

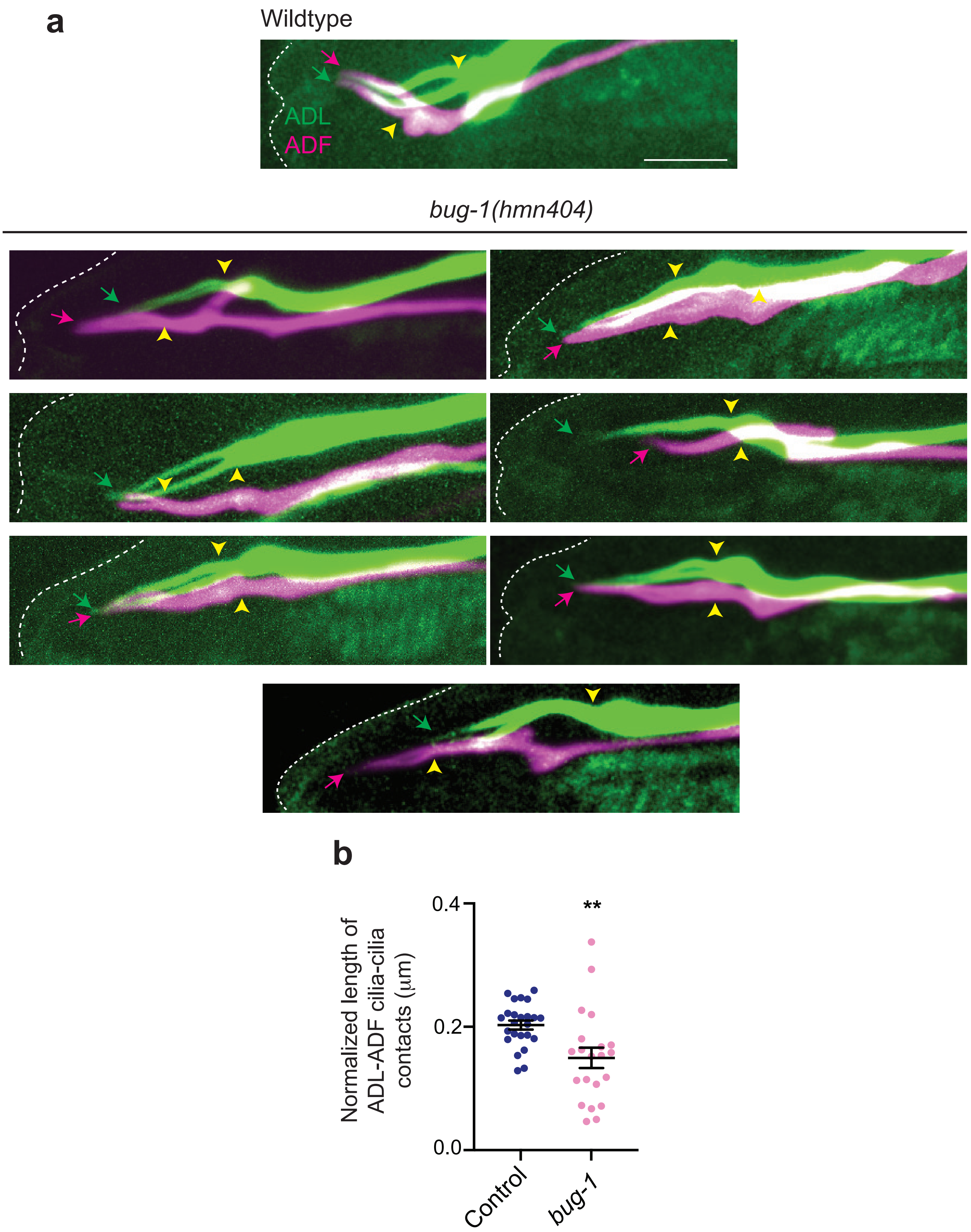
a
Wildtype
ADL
ADF
bug-1(hmn404)
b
Normalized length of ADL-ADF cilia-cilia contacts (µm)
0.4
0.2
0.0
**
Control
bug-1

**Supplementary Fig. 4. A subset of ADF and ADL cilia exhibits distinctive morphologies in *bug-1* mutants.**

**a)** Representative maximum intensity projection images of ADL and ADF cilia showing distinct cilia morphologies and interciliary contact phenotypes in *bug-1* mutants. Dashed white lines: worm nose/head; arrows: cilia tip; yellow arrowheads: cilia base. Anterior at left. Scale bar (for all images): 5 μm.

**b)** Extent of contacts between ADF and ADL cilia normalized for cilia length. Each circle is the measurement from a single pair of neurons. Data repeated from Figure 5g excluding neuron pairs that do not exhibit contacts. **: different from wildtype at $P<0.01$ (Mann-Whitney test). $n>21$ neuron pairs; three biologically independent experiments.

**Supplementary Video 1. The cilia of the ADL and ADF neurons exhibit a stereotypical contact pattern.**

(Left) Representative video of a high-resolution laser scanning confocal micrograph of ADL (green) and ADF (red) cilia in the head of an adult hermaphrodite animal. (Right) Three-dimensional rendering of the confocal image using IMARIS software (Oxford Instruments). Anterior is at left.

**Supplementary Table 1.** Strains used this work.

| Strain | Genotype | Source |
|---|---|---|
| PY13100 | *oyIs99 [srh-142*p*::dsRed, srh220*p*::gfp]* | This work |
| PY13151 | *sax-7(eq1); oyIs52 [srh-142*p*::dsRed]; oyIs57 [srh-234*p*::gfp]* | This work |
| PY13152 | *aman-2(tm1078); oyIs52 [srh-142*p*::dsRed]; oyIs57 [srh-234*p*::gfp]* | This work |
| PY13153 | *oyIs52 [srh-142*p*::dsRed]; oyIs57 [srh-234*p*::gfp]* | This work |
| PY13154 | *bbs-7(ok1351); oyIs99 [srh-142*p*::dsRed, srh-220*p*::gfp]* | This work |
| PY13155 | *kap-1(ok676); osm-3(p802); oyIs99 [srh-142*p*::dsRed, srh-220*p*::gfp]* | This work |
| PY13156 | *kap-1(ok676); osm-3(p802); oyIs99 [srh-142*p*::dsRed, srh-220*p*::gfp]; oyEx814 [srh220*p*::osm-3(oy156ts), srh-142*p*::osm-3(oy156ts)]* | This work |
| PY13157 | *kap-1(ok676); osm-3(p802); oyIs99 [srh-142*p*::dsRed, srh-220*p*::gfp]; oyEx825 [srh-142*p*::osm-3(oy156ts)]* | This work |
| PY13158 | *osm-3(p802); oyIs99 [srh-142*p*::dsRed, srh-220*p*::gfp]* | This work |
| PY13159 | *osm-3(p802); oyIs99 [srh-142*p*::dsRed, srh-220*p*::gfp]; oyEx815 [srh220*p*::osm-3(oy156ts); srh-142*p*::osm-3(oy156ts)]* line 1 | This work |
| PY13160 | *osm-3(p802); oyIs99 [srh-142*p*::dsRed, srh-220*p*::gfp]; oyEx816 [srh220*p*::osm-3(oy156ts); srh-142*p*::osm-3(oy156ts)]* line 2 | This work |
| VC1104 | *arl-13(gk513)* | CGC |
| PY13162 | *arl-13 (gk513); oyIs99 [srh-142*p*::dsRed, srh-220*p*::gfp]* | This work |
| PY13163 | *oyIs14 [sra-6*p*::gfp]; oyEx823 [flp-6*p*::mNeptune2.5]* | This work |
| PY13164 | *kap-1(ok676); osm-3(p802); oyIs14 [sra-6*p*::gfp]; oyEx818 [flp-6*p*::mNeptune2.5]* | This work |
| PY13165 | *kap-1(ok676); osm-3(p802); oyIs14 [sra-6*p*::gfp]; oyEx819 [flp-6*p*::mNeptune2.5, flp-6*p*::osm-3(oy156ts); sra-6*p*::osm-3 (oy156ts)]* line 1 | This work |

| | | |
|---|---|---|
| PY13166 | *kap-1(ok676); osm-3(p802); oyIs14 [sra-6*p*::gfp]; oyEx824 [flp-6*p*::mNeptune2.5, flp-6*p*::osm-3(oy156ts); sra-6*p*::osm-3 (oy156ts)]* line 2 | This work |
| PY13167 | *oyIs14 [sra-6*p*::gfp]; oyEx820 [srg-47*p*::mCherry]* | This work |
| PY13168 | *kap-1(ok676); osm-3(p802); oyIs14 [sra-6*p*::gfp]; oyEx821 [srg-47*p*::mCherry]* | This work |
| VC2109 | *inpp-1(gk3262)* | CGC |
| PY13170 | *inpp-1(gk3262); oyIs99 [srh-142*p*::dsRed, srh-220*p*::gfp]* | This work |
| PY13101 | *bug-1(hmn404); oyIs99 [srh-142*p*::dsRed, srh-220*p*::gfp]* | This work |
| PY12013 | *kap-1(ok676); osm-3(p802)* | (Philbrook et al. 2024) |
| PR802 | *osm-3(p802)* | CGC |
| RB1268 | *bbs-7(ok1351)* | CGC |
| CHB5782 | *bug-1(hmn404)* | (Wexler *et al.* 2026) |
| LH81 | *sax-7(eq1)* | CGC |
| FX01070 | *aman-2(tm1078)* | CGC |
| PY13173 | *kap-1(ok676); osm-3(p802); oyIs14 [sra-6*p*::gfp]; oyEx828 [srg-47*p*::osm-3(oy156ts); srg-47*p*::mNeptune2.5]* | This work |
| PY13174 | *kap-1(ok676); osm-3(p802); oyIs14 [sra-6*p*::gfp]; oyEx829 [flp-6*p*::osm-3(oy156ts); flp-6p::mNeptune2.5]* | This work |
| PY13111 | *bug-1(hmn467[bug-1::sfgfp]); oyEx790 [sra-6*p*::TagRFP; sra-6*p*::TIR1::SL2::Scarlet]* | This work, (Wexler *et al.* 2026) |
| PY13105 | *bug-1(hmn467[bug-1::sfgfp]); oyEx787 [srh-142*p*::dsRed; srh-220*p*::dsRed]* | This work, (Wexler *et al.* 2026) |
| PY13169 | *kap-1(ok676); osm-3(p802); oyIs14[sra-6*p*::gfp]; oyEx822[srg-47*p*::mNeptune2.5, sra-6*p*::osm-3(oy156ts)]* | This work |
| PY13175 | *bug-1(hmn467[bug-1::sfgfp]); oyEx837[srh-220*p*::dsRed]* | This work |

| | | |
|---|---|---|
| PY13176 | *bug-1(hmn467[bug-1::sfgfp]); oyEx838[srh-142*p*::dsRed]* | This work |
| PY13177 | *osm-3(p802); oyIs14[sra-6*p*::gfp]; oyEx839[flp-6*p*::mNeptune2.5]* | This work |
| PY13178 | *osm-3(p802); oyIs14[sra-6*p*::gfp]; oyEx839[flp-6*p*::mNeptune2.5]; oyEx840[srh-142*p*::osm-3(oy156ts); srh-220*p*::osm-3(oy156ts)]* Line 1 | This work |
| PY13179 | *osm-3(p802); oyIs14[sra-6*p*::gfp]; oyEx839[flp-6*p*::mNeptune2.5]; oyEx841[srh-142*p*::osm-3(oy156ts); srh-220*p*::osm-3(oy156ts)]* Line 2 | This work |
| PY13180 | *kap-1(ok676); osm-3(p802); oyIs99[srh-142*p*::dsRed, srh220*p*::gfp]; oyEx842[flp-6*p*::osm-3(oy156ts); sra-6*p*::osm-3(oy156ts)]* Line 1 | This work |
| PY13181 | *kap-1(ok676); osm-3(p802); oyIs99[srh-142*p*::dsRed, srh220*p*::gfp]; oyEx843[flp-6*p*::osm-3(oy156ts); sra-6*p*::osm-3(oy156ts)]* Line 2 | This work |
| PY12666 | *kap-1(ok676); osm-3(oy156ts); oyIs99[srh-142*p*::dsRed; srh-220*p*::gfp]* | This work |

**Supplementary Table 2.** Plasmids used in this work.

| Plasmid | Description | Source |
|---|---|---|
| PSAB1412 | *srh220*p*::osm-3(oy156ts)* | This work |
| PSAB1413 | *srh142*p*::osm-3(oy156ts)* | This work |
| PSAB1021 | *srh-142*p*::dsRed* | (Kazatskaya et al. 2017) |
| PSAB1419 | *srh220*p*::gfp* | This work |
| PSAB1418 | *srg-47*p*::mCherry* | (Cornils et al. 2016) |
| PSAB1415 | *srg-47*p*::mNeptune2.5* | This work |
| PSAB1414 | *sra-6p::osm-3(oy156ts)* | This work |
| PSAB1416 | *flp-6*p*::mNeptune2.5* | This work |
| PSAB1417 | *flp-6*p*::osm-3(oy156ts)* | This work |
| PSAB1343 | *sra-6p::myrTagRFP* | (Philbrook et al. 2024) |
| PSAB1422 | *sra-6p::TIR1::SL2::Scarlet* | Alison Philbrook |
| PSAB1423 | *srh-220p::dsRed* | This work |

### REFERENCES


Cornils A, Maurya AK, Tereshko L, Kennedy J, Brear AG, Prahlad V, Blacque OE, Sengupta P. 2016. Structural and functional recovery of sensory cilia in *C. elegans ift* mutants upon aging. PLoS Genet. 12:e1006325.

Kazatskaya A, Kuhns S, Lambacher NJ, Kennedy JE, Brear AG, McManus GJ, Sengupta P, Blacque OE. 2017. Primary cilium formation and ciliary protein trafficking is regulated by the atypical map kinase *mapk15* in *Caenorhabditis elegans* and human cells. Genetics. 207:1423-1440.

Philbrook A, O'Donnell MP, Grunenkovaite L, Sengupta P. 2024. Cilia structure and intraflagellar transport differentially regulate sensory response dynamics within and between *C. elegans* chemosensory neurons. PLoS Biol. 22:e3002892.
